# Analgesic α-conotoxins modulate GIRK1/2 channels via GABA_B_ receptor activation and reduce neuroexcitability

**DOI:** 10.1101/2020.12.02.407627

**Authors:** Anuja R. Bony, Jeffrey R. McArthur, Rocio K. Finol-Urdaneta, David J. Adams

## Abstract

Activation of G protein-coupled inwardly rectifying potassium (GIRK or Kir3) channels leads to membrane hyperpolarization and dampening of neuronal excitability. Here we show that the analgesic α-conotoxin Vc1.1 potentiates inwardly rectifying K^+^ currents (I_Kir_) mediated through native and recombinant GIRK1/2 channels by activation of the G protein-coupled GABA_B_ receptor (GABA_B_R) via a *Pertussis* toxin (PTX)-sensitive G protein. Recombinant co-expression of human GIRK1/2 subunits and GABA_B_R in HEK293T cells resulted in a Ba^2+^-sensitive I_Kir_ potentiated by baclofen and Vc1.1 which was inhibited by PTX, intracellular GDP-β-S, or the GABA_B_R-selective antagonist CGP 55845. In adult mouse DRG neurons, GABA_B_R-dependent GIRK channel potentiation by Vc1.1 and baclofen hyperpolarizes the cell resting membrane potential with concomitant reduction of excitability consistent with Vc1.1 and baclofen analgesic effects *in vivo*. This study provides new insight into Vc1.1 as an allosteric agonist for GABA_B_R-mediated potentiation of GIRK channels and may aid in the development of novel non-opioid treatments for chronic pain.

## Introduction

Current pharmacotherapies available for the treatment of neuropathic and chronic pain include opioids and anti-depressants frequently associated with severe adverse effects (Costigan *et al*, 2009; Finnerup *et al*, 2021) underscoring the need for alternative therapies. Several venom-derived peptides have been demonstrated to selectively target neurotransmitter receptors and ion channels in nociceptive pathways. In particular, a number of relatively small disulfide-rich peptides isolated from marine cone snails called conotoxins exhibit anti-nociceptive activity (Knapp *et al*, 2012; Sadeghi *et al*, 2017; Hone *et al*, 2018; Kennedy *et al*, 2020). ω-Conotoxin MVIIA, which inhibits neuronal high voltage-activated (HVA) N-type (Cav2.2) calcium channels, is the active element of Prialt™ (Ziconotide), an FDA approved drug for the treatment of intractable pain (Wermeling, 2005). However, off-target effects and the requirement for intrathecal administration restrict its applications.

α-Conotoxins are primarily recognised as muscle and neuronal nicotinic acetylcholine receptor (nAChR) antagonists (Lebbe *et al*, 2014; Abraham & Lewis, 2018). Yet, α-conotoxins active at α9α10 nAChRs also inhibit neuronal calcium (Cav) channels via activation of G protein-coupled GABA_B_ receptors (GABA_B_Rs) (Callaghan *et al*, 2008; Daly *et al*, 2011; Adams *et al*, 2012; Cuny *et al*, 2012). α-Conotoxin Vc1.1 from *Conus victoriae* venom was shown to antagonize α9α10 nAChRs (Vincler *et al*, 2006; Nevin *et al*, 2007) and to activate GABA_B_Rs resulting in inhibition of Cav2.2 (N-type) and Cav2.3 (R-type) channels in mammalian primary afferent neurons (Callaghan *et al*, 2008; Berecki *et al*, 2014; Sadeghi *et al*, 2017). In α9 knockout mice and rodent neuropathic and chronic visceral pain models, Vc1.1 and its analogues, exhibit anti-nociceptive activity via GABA_B_R inhibition of Cav2.2/Cav2.3-mediated currents (Callaghan & Adams, 2010; Mohammadi & Christie, 2014; Castro *et al*, 2018). Selective GABA_B_R antagonists, and Gα_i/o_ inhibitors such as *Pertussis* toxin, GDP-β-S, and G protein βγ scavengers, all attenuate α-conotoxin inhibition of Cav2.2/Cav2.3 channels. In HEK293 cells co-expressing Cav2.2 and GABA_B_R where the GABA/baclofen binding sites have been neutralized, Vc1.1 remains able to inhibit Cav2.2 Ca^2+^ currents suggesting that it acts as an allosteric GABA_B_R agonist (Huynh *et al*, 2015). The cyclized Vc1.1 analogue, cVc1.1, is >8000-fold more potent inhibiting GABA_B_R-dependent HVA calcium channels than α9α10 nAChRs highlighting the role of GABA_B_R in its anti-nociceptive actions (Clark *et al*, 2010; Castro *et al*, 2018; Sadeghi *et al*, 2018).

GABA_B_Rs functionally couple to GIRK channels to attenuate nociceptive transmission (Blednov *et al*, 2003) analogous to the analgesic effects observed upon agonist activation of μ-opioid, α2 adrenergic, muscarinic or cannabinoid receptors (Blednov *et al*, 2003; Mitrovic *et al*, 2003; Nagi & Pineyro, 2014; Yudin & Rohacs, 2018), or by direct activation of GIRK channels (Lujan *et al*, 2014). GIRK channels are tetrameric assemblies of four distinct subunits, GIRK1, GIRK2, GIRK3 and GIRK4, of which only GIRK2 and GIRK4 channels form functional homotetramers whereas GIRK1 and GIRK3 are obligatory heterotetramers requiring other GIRK subunits to enable trafficking to the plasma membrane (Lujan *et al*, 2014; Lujan *et al*, 2015). All GIRK subunits, but in particular GIRK1 and GIRK2, are expressed in mammalian sensory neurons where they may couple with G protein-coupled receptors (GPCRs) including GABA_B_R (Marker *et al*, 2004; Kanjhan *et al*, 2005; Gao *et al*, 2007; Lyu *et al*, 2015). Direct binding of the G protein (G_i/o_) Gβγ subunit to the GIRK channel has been shown to activate and modulate the inhibitory actions of several neurotransmitters (Hibino *et al*, 2010; Luscher & Slesinger, 2010;). Independent of GPCR activation, a number of clinically active compounds targeting GIRK channels have been described to modulate diverse pathological processes (Luscher & Slesinger, 2010; Wydeven *et al*, 2014). For example, the GIRK channel opener VU046655 exhibits anti-nociceptive activity implicating GIRK-mediated activity in analgesia (Abney *et al*, 2019).

In the present study, we investigated the effect of the analgesic α-conotoxins Vc1.1, RgIA, and PeIA on recombinant and native GIRK-mediated K^+^ currents and their effects on neuronal excitability. These conopeptides potentiated heteromeric GIRK1/2 and homomeric GIRK2 channels when co-expressed with human GABA_B_R in HEK293T cells and supressed excitability in adult mouse DRG neurons. A preliminary account of the results has been presented as a published abstract (Bony *et al*, 2020).

## Results

### Analgesic α-conotoxins potentiate GIRK1/2 channel-mediated K^+^ currents via GABA_B_R activation

The effect of α-conotoxin Vc1.1 on GIRK1/2 channels was investigated in HEK293T cells transiently transfected with human GIRK1, GIRK2, GABA_B_R1 and GABA_B_R2 subunits. In transfected HEK293T cells, basal inwardly rectifying K^+^ current (I_Kir_) density recorded in the presence of 20 mM extracellular K^+^ at −100 mV was 55.7 ± 7.5 pA/pF (n = 17). I_Kir_ was reversibly reduced by ~75% to 18.9 ± 4.1 pA/pF (n = 17) upon bath application of 1 mM Ba^2+^ which effectively blocks most Kir channels (Hibino *et al*, 2010). Under our recording conditions, non-transfected HEK293 cells exhibited an endogenous I_Kir_ density of 1.5 ± 0.2 pA/pF (n = 6) consistent with that reported previously (Ämmälä *et al*, 1996). Bath application of either Vc1.1 (1 μM) or baclofen (100 μM) reversibly potentiated whole-cell inward K^+^ currents at −100 mV by 28.3 ± 1.8 pA/pF (n = 8) and 68.4 ± 5 pA/pF (n = 9), respectively (Fig. 1A, B; Table 1). Barium (1 mM) fully blocked the potentiation of I_Kir_ by Vc1.1 and baclofen (Fig. 1A, B). Vc1.1 and baclofen were ineffective at potentiating GIRK1/2-mediated I_Kir_ in HEK293 cells expressing GIRK1/2 channels alone in the absence of GABA_B_R (n = 5; Fig. 1C, D & H). Since GIRK2 can form functional homotetramers, the effect of α-conotoxin Vc1.1 was verified on cells co-expressing homomeric GIRK2 channels and GABA_B_Rs. The basal I_Kir_ density for GIRK2 transfected cells was 17.3 ± 4.4 pA/pF (n = 12) which was reversibly inhibited by >63% in the presence of 1 mM Ba^2+^. Vc1.1 (1 μM) and baclofen (100 μM) reversibly potentiated GIRK2-mediated I_Kir_ at −100 mV by 9.8 ± 1.0 pA/pF (n = 6) and 27.1 ± 4.2 pA/pF (n = 4), respectively (Fig. 1E, F & H; Table 1).

**Figure 1.**
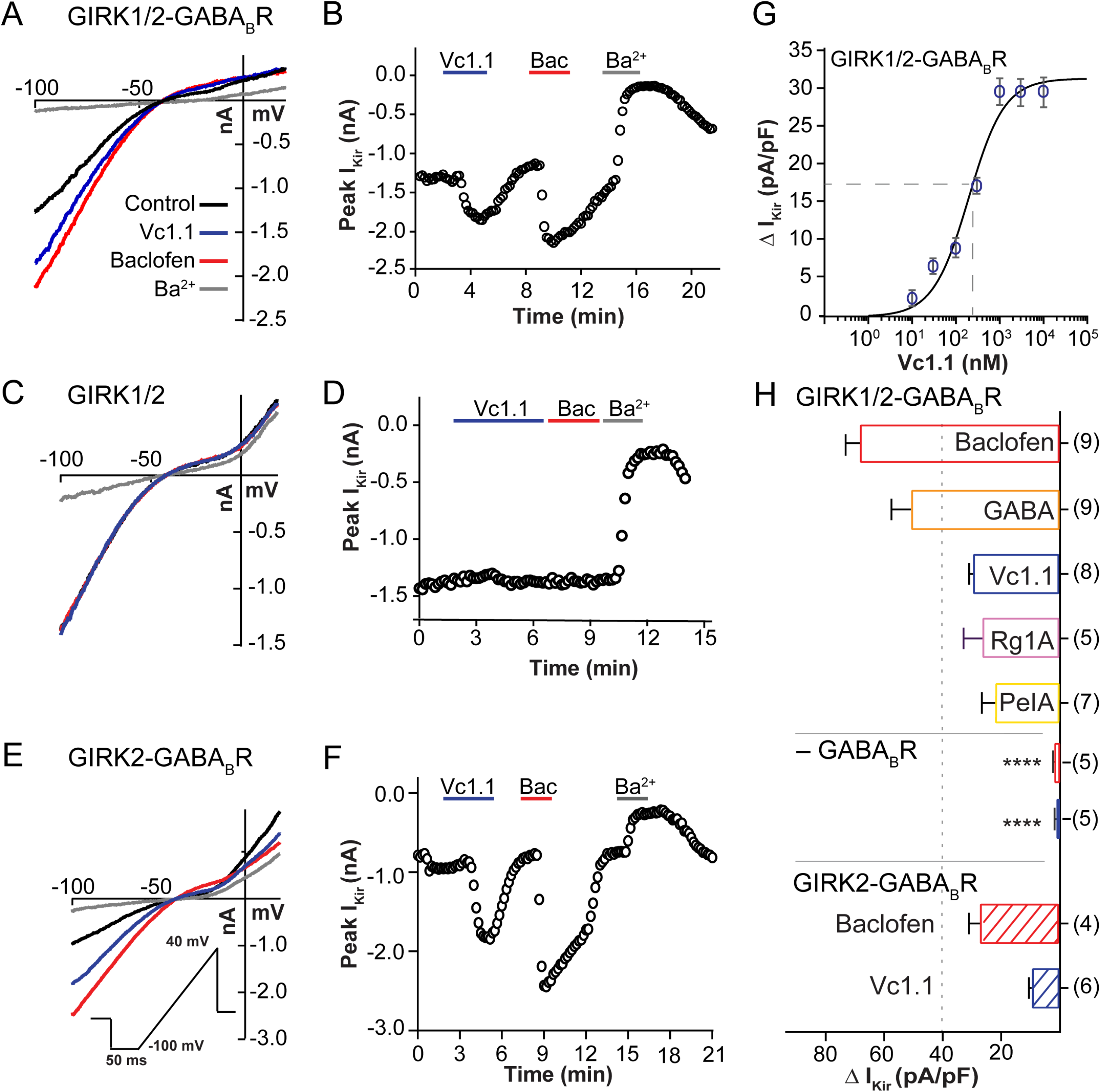
α-Conotoxin Vc1.1 modulation of heteromeric GIRK1,2 and homomeric GIRK2 channels require GABA_B_R. **A, C** Representative K^+^ currents recorded from GIRK1/2 channels co-expressed either with GABA_B_R (**A**) or alone (**C**) in response to a 50 ms voltage ramp protocol (−100 to +40 mV; see inset) applied at 0.1 Hz from a holding potential of −40 mV in the absence (control) and presence of 1 μM Vc1.1 (blue), 100 μM baclofen (red) or 1 mM Ba^2+^ (grey). **B, D** Corresponding diary plots of K^+^ current amplitude at −100 mV as a function of time in cells expressing GIRK1/2 channels and GABA_B_R **(B)** and GIRK1/2 channels alone **D** Responses to sequential bath application of Vc1.1 (blue), baclofen (red), and Ba^2+^ (grey) are indicated by the bars above. **E** Vc1.1 potentiates also homomeric GIRK2 channel when co-expressed with GABA_B_R. Representative human GIRK2-mediated K^+^ currents obtained in the absence (Control, black), and presence of 1 μM Vc1.1 (blue), 100 μM baclofen (red) or 1 mM Ba^2+^ (grey). **F** Corresponding diary plot to (E) showing peak K^+^ current amplitude at −100 mV as a function of time in response to bath application of Vc1.1 (blue), baclofen (red) or Ba^2+^ (grey). **G** α-Conotoxin Vc1.1 concentration-response relationship obtained for potentiation of GIRK1/2 co-expressed with GABA_B_R in HEK293T cells. ΔI_Kir_ represents total I_Kir_ (basal + potentiation) minus basal I_Kir_. **H** Bar graph of ΔI_Kir_ density (pA/pF) at −100 mV in response to 100 μM baclofen (red), 100 μM GABA (orange), 1 μM Vc1.1 (blue), 1 μM RgIA (purple) and 1 μM PeIA (yellow) recorded from cells expressing either heteromeric GIRK1/2 (solid) or homomeric GIRK2 (hashed) co-expressed with GABA_B_R. Neither Vc1.1 nor baclofen potentiates K^+^ currents in cells expressing GIRK1/2 alone. The vertical dotted line (40 pA/pF) is for reference. Data represent mean ± SEM and the number of experiments is given in parentheses.

**Table 1.**
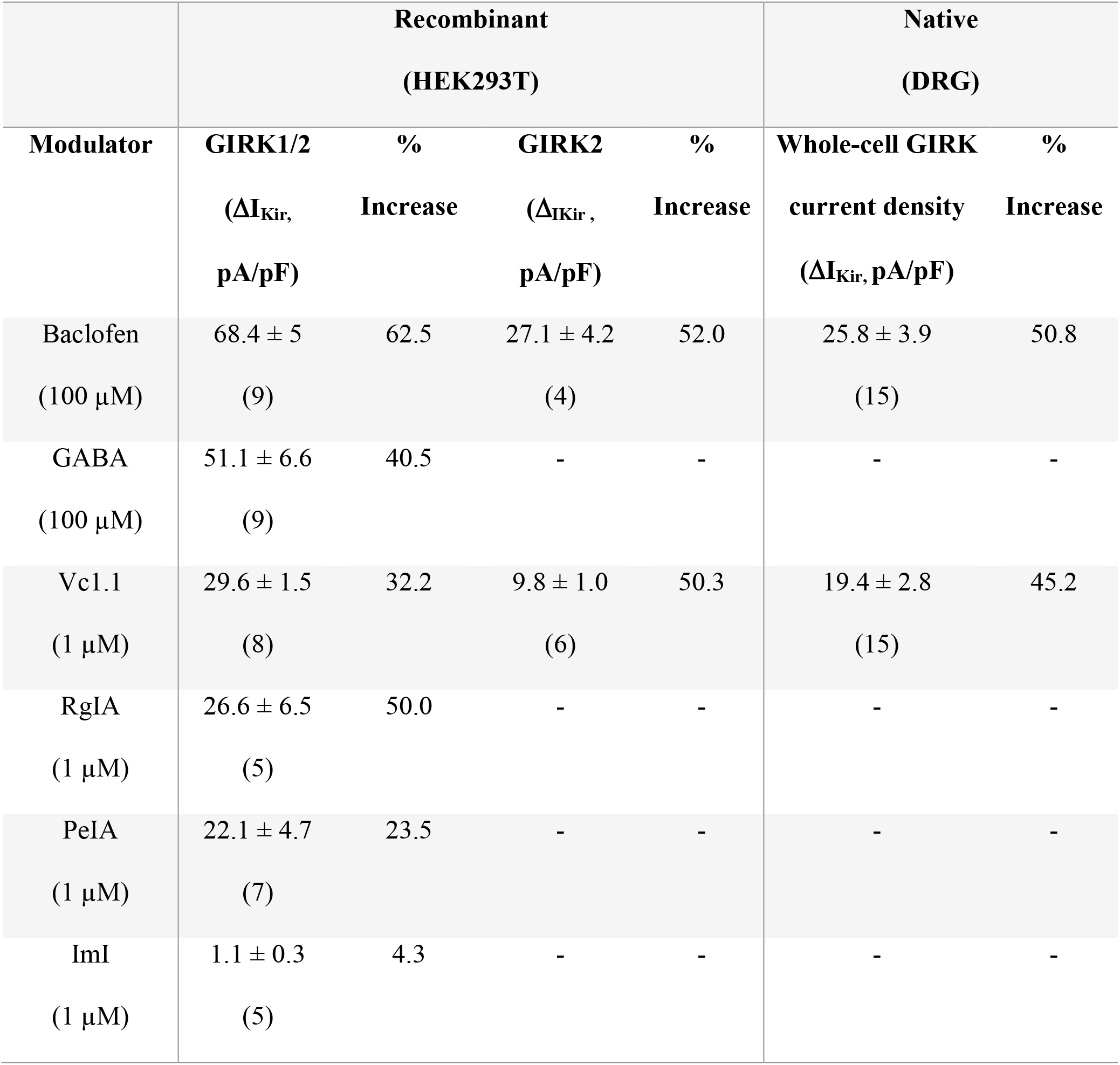
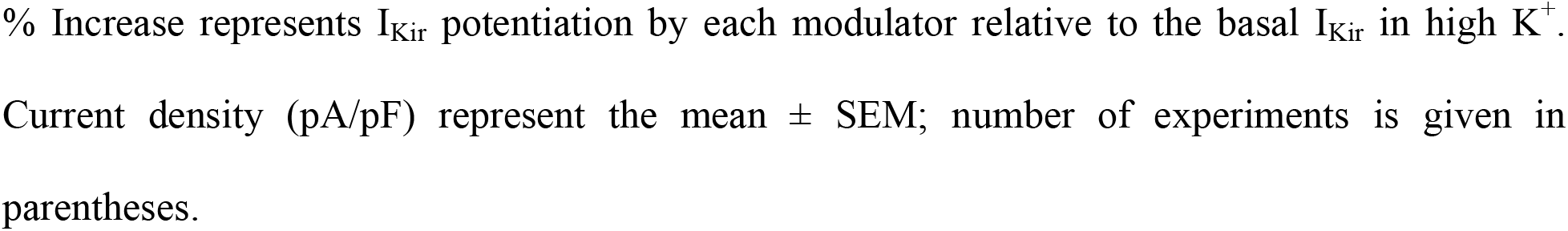
Summary of GIRK-mediated K^+^ current potentiation (ΔI_Kir_) at −100 mV by Baclofen, GABA, and α-conotoxins Vc1.1. RgIA, PeIA and ImI in HEK293T cells co-expressing GABA_B_R and either GIRK1/2 or GIRK2 channels and mouse DRG neurons.

| Modulator | Recombinant<br>(HEK293T) |  |  |  | Native<br>(DRG) |  |
| --- | --- | --- | --- | --- | --- | --- |
| | GIRK1/2<br>($\Delta I_{Kir}$,<br>pA/pF) | %<br>Increase | GIRK2<br>($\Delta I_{Kir}$,<br>pA/pF) | %<br>Increase | Whole-cell GIRK<br>current density<br>($\Delta I_{Kir}$, pA/pF) | %<br>Increase |
| Baclofen<br>(100 μM) | 68.4 ± 5<br>(9) | 62.5 | 27.1 ± 4.2<br>(4) | 52.0 | 25.8 ± 3.9<br>(15) | 50.8 |
| GABA<br>(100 μM) | 51.1 ± 6.6<br>(9) | 40.5 | - | - | - | - |
| Vc1.1<br>(1 μM) | 29.6 ± 1.5<br>(8) | 32.2 | 9.8 ± 1.0<br>(6) | 50.3 | 19.4 ± 2.8<br>(15) | 45.2 |
| RgIA<br>(1 μM) | 26.6 ± 6.5<br>(5) | 50.0 | - | - | - | - |
| PeIA<br>(1 μM) | 22.1 ± 4.7<br>(7) | 23.5 | - | - | - | - |
| ImI<br>(1 μM) | 1.1 ± 0.3<br>(5) | 4.3 | - | - | - | - |

% Increase represents $I_{Kir}$ potentiation by each modulator relative to the basal $I_{Kir}$ in high $K^+$ . Current density (pA/pF) represent the mean ± SEM; number of experiments is given in parentheses.

Other α-conotoxins active at α9α10 nAChRs including RgIA and PeIA potently inhibit HVA calcium channels via GABA_B_R activation (Callaghan *et al*, 2008; Daly *et al*, 2011; Berecki *et al*, 2014) and therefore were also tested on recombinant GIRK1/2-GABA_B_R in HEK293T cells. Bath application of α-conotoxins PeIA and RgIA at 1 μM reversibly potentiated the GIRK1/2-mediated I_Kir_ increasing the current density by 26.6 ± 6.5 pA/pF and 22.1 ± 4.7 pA/pF (n = 5-7), respectively (Fig. 1H; Table 1). In contrast, application of the α7 nAChR antagonist, α-conotoxin ImI (1 μM), failed to potentiate I_Kir_ (n = 5) compared to Vc1.1 tested in the same cells (Fig. EV1C, D & E). Furthermore, reduction of Vc1.1 (Vc1.1^RA^) with dithiothreitol (DTT; 1 mM) rendered the peptide incapable of potentiating GIRK1/2-mediated I_Kir_ (n = 5; Fig. EV1A, B & E).

Vc1.1 potently potentiated GABA_B_R-coupled GIRK1/2 channels with an EC_50_ of 197.2 ± 0.2 nM (n^H^ = 1.1; n = 5-7 replicates per concentration) determined from the concentration-response relationship displayed in Fig. 1G. Notably, the concentration of Vc1.1 to potentiate GABA_B_R-GIRK1/2 K^+^ currents is ~100-fold higher than that required for Vc1.1 inhibition of rodent DRG neuron GABA_B_R-HVA calcium currents and recombinant human GABA_B_R-Cav2.2 expressed in HEK293 cells (Callaghan *et al*, 2008; Huynh *et al*, 2015).

### GIRK1/2 channels are not directly activated by Vc1.1

The involvement of GABA_B_R in the potentiation of GIRK1/2 and GIRK2 K^+^ currents by analgesic α-conotoxins and baclofen was investigated in HEK293T cells by expressing GIRK1/2 alone and using the selective GABA_B_R antagonist, CGP 55845 upon co-transfection with GABA_B_R. In the absence of GABA_B_R expression, application of either baclofen or Vc1.1 failed to potentiate GIRK1/2 K^+^ currents indicating that Vc1.1 and baclofen potentiation of recombinant I_Kir_ is GABA_B_R-dependent. Bath application of CGP 55845 (1 μM) did not affect the GIRK1/2 K^+^ current but robustly and reversibly antagonized the potentiation of I_Kir_ by both Vc1.1 (80%, n = 5) and baclofen (88%, n = 5), respectively (Fig. 2A, B & I). In HEK293T cells, CGP 55845 antagonism of Vc1.1- and baclofen-induced potentiation of I_Kir_ was reversible. Taken together, these results highlight a requirement for GABA_B_R activation for the potentiation of GIRK1/2 K^+^ currents by Vc1.1 and baclofen.

**Figure 2.**
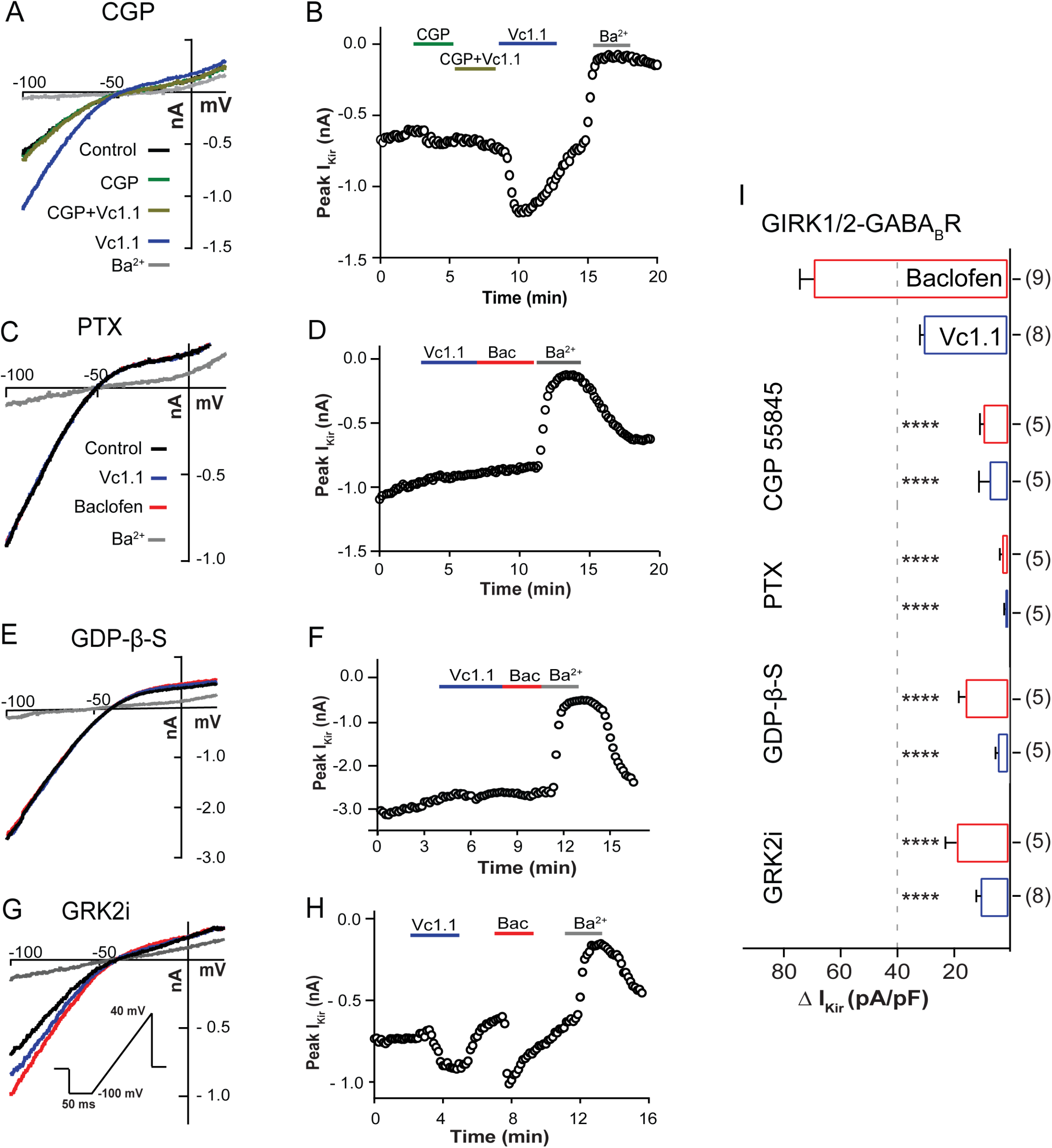
Potentiation of GIRK1/2-mediated K^+^ current by either α-conotoxin Vc1.1 or baclofen requires co-expression of GABA_B_R and involves G protein Gα_i/o_ and βγ. **A** Bath application of the selective GABA_B_R antagonist CGP 55845 (1 μM) antagonizes Vc1.1 potentiation of GIRK1/2 channels. Representative K^+^ currents recorded from HEK293T cells co-expressing GABA_B_R and GIRK 1/2 channels in the absence (control; black) or presence of 1 μM CGP 55845 alone (green), and co-application of CGP 55845 plus 1 μM Vc1.1 (light green), Vc1.1 alone (blue) or 1 mM Ba^2+^ (grey). **B** Corresponding dairy plot of K^+^ current amplitude at −100 mV (A) as a function of time. **C** Representative K^+^ currents elicited from cells co-expressing GABA_B_R and GIRK1/2 channels with 24 hr pre-treatment with 1 μg/ml *Pertussis* toxin (PTX). **D** Corresponding diary plots of peak K^+^ currents at −100 mV (C) as a function of time. **E** Inclusion of 500 μM GDP-β-S in the intracellular patch pipette solution abolished Vc1.1 potentiation of GIRK1/2 current. Representative K^+^ current traces in the absence (control; black) or presence of Vc1.1 (blue), baclofen (red) or Ba^2+^ (grey). **F** Corresponding diary plot of (E) as a function of time. **G** Inclusion of Gβγ scavenger, GRK2i (10 μM) inhibited potentiation by both Vc1.1 and baclofen in cells co-expressing GABA_B_R and GIRK1/2. **H** Corresponding time plot of (G) showing peak I_Kir_ potentiation by Vc1.1 and baclofen at −100 mV. **I** Bar graph of ΔI_Kir_ density recorded at −100 mV in response to 1 μM Vc1.1 (−29.5 ± 1.52 pA/pF, blue) or 100 μM baclofen (−68.4 ± 5 pA/pF, red) in HEK293T cells co-expressing GIRK1/2 channels and GABA_B_R (control). Potentiation of ΔI_Kir_ density by Vc1.1 and baclofen was significantly reduced by 1 μM CGP 55845. Following pre-treatment with PTX (1 μg/ml) or inclusion of either GDP-β-S (500 μM) or Gβγ scavenger, GRK2i (10 μM), in the intracellular pipette solution, ΔI_Kir_ density was similarly attenuated compared to peak current density measured for Vc1.1 or baclofen in control condition. Data are expressed as mean ± SEM and statistical significance, **** p < 0.0001 vs control; one-way ANOVA followed by Tukey’s post hoc test; number of experiments is given in parentheses.

### Gβγ mediates Vc1.1 modulation of GIRK1/2

GIRK channel modulators like the orthosteric agonists, GABA and baclofen, act via the G protein Gα_i/o_, whereas Gβγ is known to interact with the GIRK1 and GIRK2 N-terminus or C-terminus (Ivanina *et al*, 2003; Kahanovitch *et al*, 2014; Dascal & Kahanovitch, 2015). The role of G proteins in GABA_B_R-mediated potentiation of GIRK channels by Vc1.1 was investigated using selective G protein inhibitors. A 24 hour incubation with *Pertussis* toxin (PTX; 1 μg/ml), an inhibitor of heterotrimeric G_i/o_ proteins (Gα_i/o_), abrogated the potentiation of GIRK1/2-mediated K^+^ currents by 1 μM Vc1.1 to 1.1 ± 0.5 pA/pF (n = 5) and 100 μM baclofen to 2.0 ± 0.8 pA/pF (n = 5) compared to untreated cells (Fig. 2C, D & I). Replacement of GTP with the hydrolysis-resistant GDP analogue, GDP-β-S (500 μM) in the patch pipette, attenuated Vc1.1 and baclofen potentiation of GABA_B_R-GIRK1/2 K^+^ currents compared to control (3.6 ± 0.9 pA/pF (n = 5) and 15.4 ± 2.3 pA/pF (n = 5), respectively) (Fig. 2E, F & I). In contrast, inclusion of the non-hydrolyzable G protein-activating GTP analogue, GTP-γ-S (500 μM) in the patch pipette, potentiated basal I_Kir_ approximately two-fold from control (55.7 ± 7.5 pA/pF) to GTP-γ-S (114.8 ± 9.4 pA/pF, n = 5; p = 0.0004) whereas Vc1.1 (1 μM) potentiation of I_Kir_ was increased by 78.1% in the presence of GTP-γ-S (from 94.6 ± 8.7 pA/pF to 168.5 ± 4.0 pA/pF, n = 5; p = 0.0051) (Fig. EV2A, B & C). Similarly, baclofen potentiation of I_Kir_ was increased 61.5% in the presence of GTP-γ-S from 114.7 ±9.4 pA/pF to 185.2 ± 20.5 pA/pF (n = 5; p = 0.002) (Fig. EV2A, B & C). The observed increase in I_Kir_ density is consistent with a contribution of Vc1.1 to the activation of Gα_i/o_ protein underlying GIRK1/2 channel potentiation. The requirement of Gβγ in the signalling cascade between the GABA_B_R and GIRK1/2 channels was investigated by supplementing the intracellular solution (patch pipette) with the peptide analogue of the G protein receptor kinase 2 (GRK2i) that acts as a Gβγ scavenger (Koch *et al*, 1994). In comparison to non-supplemented controls, GRK2i (10 μM) significantly attenuated the potentiation of GIRK1/2 K^+^ currents by Vc1.1 and baclofen by 70% and 74% (n = 5-8), respectively (Fig. 2G, H & I). Overall, these results indicate that the G protein heterodimer subunit Gβγ is involved in the modulation of GIRK1/2 channels by Vc1.1 and baclofen.

### Vc1.1 potentiates inwardly rectifying K^+^ currents and depresses excitability in mouse DRG neurons

All GIRK channel members have been identified in rat DRG neurons (Gao *et al*, 2007), however, the presence of GIRK1 and GIRK2 subunits in mouse DRG has been contested (Nockemann *et al*, 2013). The expression and co-localization of the GIRK1, GIRK2, and GABA_B_R2 proteins in mouse DRG neurons was investigated using immunofluorescence and confocal microscopy. Fig. 3A shows images of double immunostainings in which positive immunoreactivity for GIRK1 (red) and GIRK2 (green) antibodies is demonstrated. The merged panel shows that these two GIRK subunits co-localize in ~81% of the DRG neurons (13 of 16 counted) present in the imaged field. Antibody validation experiments were performed in HEK293 cells transfected with GIRK1, GIRK2 and GABA_B_R2 (Fig. EV3). Double staining was also carried out to investigate GABA_B_R and GIRK channel co-localization in mouse DRG neurons which show abundant expression of GIRK2 and GABA_B_R2 (Fig. 3B). The fluorescent signals overlapped in 96% of the DRG neurons (96 of 100 counted) indicative of co-localization (Fig. 3B merge). Due to antibody compatibility constraints, we consider immunodetection of GIRK2 and GABA_B_R2 as expression proxies, supported by previous studies showing that GIRK1 requires GIRK2 for trafficking to the plasma membrane (Hibino *et al*, 2010; Luscher & Slesinger, 2010) and that GABA_B_R can only function as R1/R2 heterodimers (Pinard *et al*, 2010; Cuny *et al*, 2012; Frangaj & Fan, 2018; Mao *et al*, 2020).

**Figure 3.**
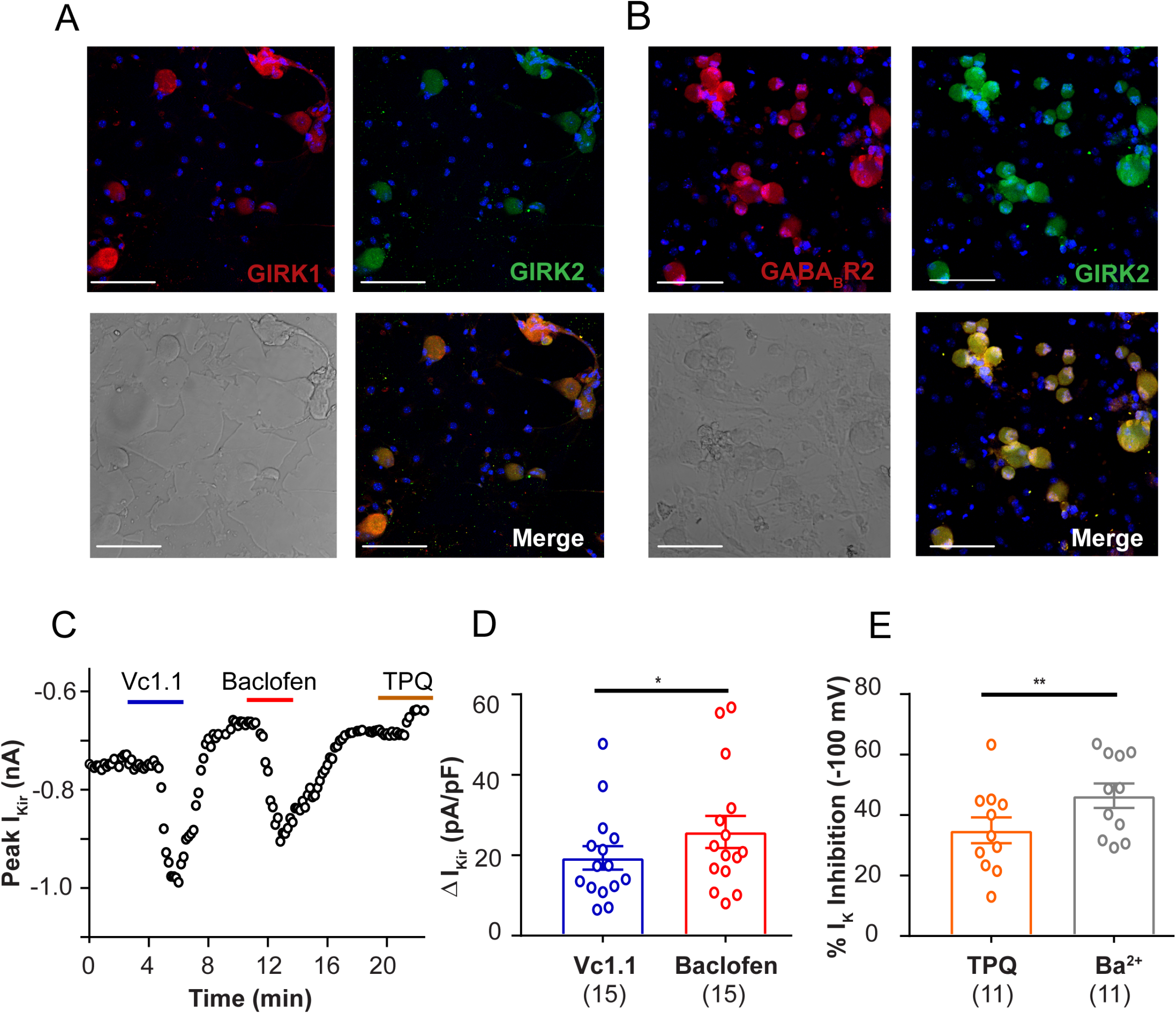
Expression of GIRK1, GIRK2 and GABA_B2_ subunit of GABA_B_R and α-conotoxin Vc1.1 potentiation of inward rectifier K^+^ currents in adult mouse DRG neurons. **A** Immunolabeling of mouse DRG neurons and visual inspection by confocal microscopy revealed both GIRK1 and GIRK2 channels express and show co-localization with each other using antibodies directed against GIRK1 and GIRK2 channels. **B** DRG neurons showing expression of GABA_B2_ subunit and GIRK2 channel and both GIRK2 and GABA_B2_ co-localize in mouse DRG neurons. Co-expression of GIRK 1 and GIRK 2 or GIRK2 and GABA_B_R immunoreactivity in mouse DRG neurons was independent of cell diameter. Scale bars: 100 μm. DAPI (blue) staining marks nucleus. **C** Diary plot of peak inward K^+^ currents recorded at −100 mV in small to medium diameter (<30 μm) mouse DRG neurons as a function of time. Whole-cell K^+^ currents obtained in response to voltage ramps (−100 mV to +40 mV) applied at 0.1 Hz in the absence (basal control taken after baclofen washout, black) and presence of 1 μM Vc1.1 (blue), 100 μM baclofen (red) or 500 nM Tertiapin-Q (brown). **D** Bar graph of inward K^+^ current density measured at −100 mV in response to Vc1.1 (1 μM, blue) and baclofen (100 μM, red). Data represent mean ± SEM. Number of experiments are shown in parentheses. p = 0.011, paired t-test. **E** Summary of the effects of Tertiapin-Q (500 nM, brown) and Ba^2+^ (1 mM, grey) on whole-cell inward I_K_. Data represent mean ± SEM. p = 0.001, paired t-test. Number of experiments is given in parentheses.

Modulation of I_Kir_ by Vc1.1 and baclofen was examined in isolated DRG sensory neurons from adult mice. Bath application of baclofen (100 μM) and Vc1.1 (1 μM) potentiated I_Kir_ elicited by 50 ms ramps from −100 mV to +40 mV in approximately one-third (15/50) of small to medium diameter (<30 μm) DRG neurons. Neuronal Ba^2+^-sensitive, inward rectifying K^+^ currents (I_Kir_) recorded in the presence of high K^+^ were potentiated by 45.2% in the presence of 1 μM Vc1.1 (ΔI_Kir_ = 19.4 ± 2.8 pA/pF; n = 15) from the control level (Fig. 3C, D; Table 1). Accordingly, baclofen (100 μM) applied to the same neurons increased I_Kir_ by 50.8% (ΔI_Kir_ = 25.8 ± 3.9 pA/pF; n = 15) consistent with GABA_B_R activation of GIRK channels in mouse DRG neurons (Fig. 3C, D; Table 1). Incubation of mouse DRG neurons with PTX (1 μg/ml) for 24 hr inhibited both Vc1.1 and baclofen potentiation of I_Kir_ (n = 5) consistent with our observation in recombinant human GABA_B_R and GIRK1/2 channels.

Inhibition of I_Kir_ by Tertiapin-Q (500 nM) was used as a reporter of GIRK channel activity (Kanjhan *et al*, 2005). Under these experimental conditions, Tertiapin-Q inhibited the whole-cell inward I_K_ recorded in high external K^+^ by 35.0 ± 4.3% whereas Ba^2+^ inhibited 46.4 ± 4.0% inward I_K_ in the same cell (n = 11, p = 0.001, paired t-test) (Fig. 3E). The co-localisation of GABA_B_R and GIRK channels in mouse DRG neurons and antagonism of Vc1.1- and baclofen-dependent potentiation of I_Kir_ by Tertiapin-Q are both consistent with the modulation of native GIRK K^+^ currents by α-conotoxin Vc1.1 and baclofen via GABA_B_R in DRG neurons.

### Vc1.1 depresses excitability in adult mouse DRG neurons

To determine if GABA_B_R-mediated GIRK channel potentiation by Vc1.1 influences neuronal excitability, small to medium diameter (<30 μm) adult mouse DRG neurons were studied under current clamp and the passive and active membrane properties compared between control and treated cells (Table 2). Bath application of α-conotoxin Vc1.1 (1 μM) reversibly reduced action potential firing in response to depolarizing current pulses in adult mouse DRG neurons (Fig. 4A). In 26 of 42 (62%) neurons recorded, the resting membrane potential was hyperpolarized by ~5 mV in the presence of 1 μM Vc1.1 (Control = −53 ± 2 mV, Vc1.1 = −58 ± 2 mV n = 24; p = 0.0024) (Fig. 4Bi, Table 2) and the input resistance (R_i_) was reduced by >1.5-fold from 542.4 ± 121.4 MΩ (control) to 344.1 ± 71.1 MΩ (n = 6; p = 0.0318) (Fig. 4Bii; Table 2). In the presence of Vc1.1, the rheobase for action potential firing increased approximately two-fold from 104 ± 18 pA (control) to 192 ± 30 pA (n = 26; p < 0.0001) (Fig. 4Biii, Table 2). Furthermore, the number of action potentials evoked in DRG neurons in response to 10 pA depolarizing current steps (−25 - 300 pA) was more than halved in the presence of Vc1.1 compared to control (n = 26; p = 0.002) (Fig. 4Biv, Table 2).

**Table 2.** Summary of the effects of α-conotoxin Vc1.1, Baclofen and Tertiapin-Q on the passive and active membrane properties of mouse DRG neurons.

| **Parameters** | **Vc1.1**<br>**(1 μM)** | **Baclofen**<br>**(100 μM)** | **Tertiapin-Q**<br>**(500 nM)** |
| --- | --- | --- | --- |
| **$\Delta$ RMP (mV)** | $-4.3 \pm 1.1$<br>(25) | $-5.1 \pm 1.9$<br>(18) | $+10.0 \pm 1.0$<br>(21) |
| **$R_i$ (fold change)** | $\downarrow 1.5 \pm 0.2$<br>(6) | $\downarrow 1.7 \pm 0.1$<br>(10) | $\uparrow 1.9 \pm 0.3$<br>(14) |
| **AP# (fold change)** | $\downarrow 1.8 \pm 0.2$<br>(26) | $\downarrow 1.5 \pm 0.3$<br>(21) | $\uparrow 3.4 \pm 0.7$<br>(21) |
| **Rheobase (fold change)** | $\uparrow 2.0 \pm 0.2$<br>(26) | $\uparrow 3.1 \pm 0.7$<br>(21) | $\downarrow 1.9 \pm 0.1$<br>(21) |

AP#, Number of action potentials; $R_i$ , Input resistance; RMP, Resting membrane potential
↑ , increase; ↓ , decrease; Fold change calculated for each individual neuron and data represent the mean ± SEM; number of experiments is given in parentheses.

**Figure 4.**
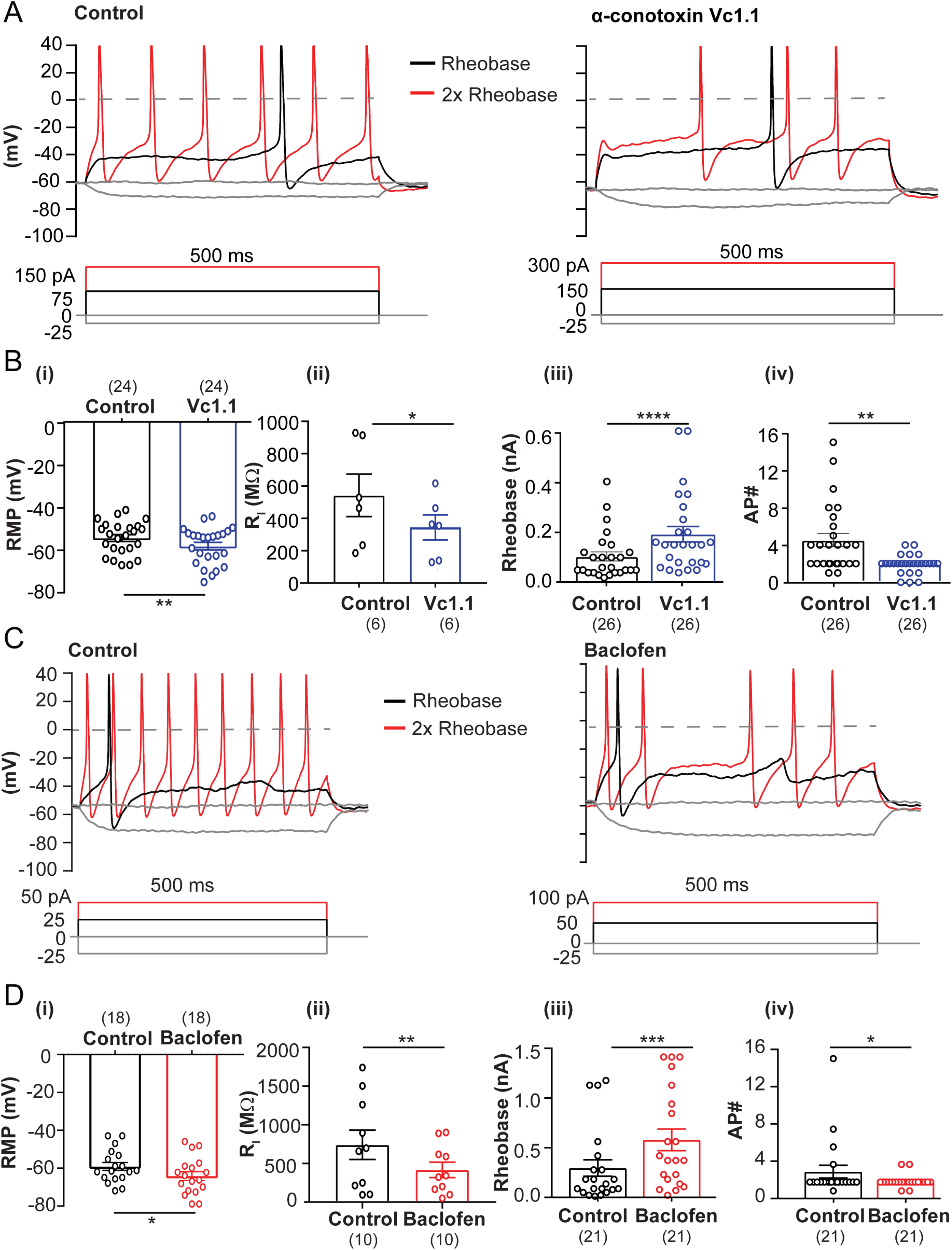
α-Conotoxin Vc1.1 hyperpolarizes the resting membrane potential and reduces excitability in mouse DRG neurons similar to the canonical GABA_B_R agonist baclofen. **A** Representative voltage responses to current clamp steps recorded in a DRG neuron (28.7 μm diameter) in the absence (control; left) and presence of 1 μM Vc1.1 (right). Broken line indicates 0 mV, black indicates the rheobase and red indicates 2x rheobase for both membrane potential and current. **B** Bar graphs and scatter plots of the effects of 1 μM Vc1.1 on resting membrane potential (RMP), p = 0.0011 **(i)**, input resistance (R_i_), p = 0.0304 **(ii)**, rheobase, p < 0.0001 **(iii)**, and number of APs, p = 0.0016 **(iv)** in response to 500 ms depolarizing current step in small to medium diameter neurons of adult mice DRG. Data represented as mean ± SEM; Paired t-test. Number of experiments are given in parentheses. **C** Representative voltage responses to current clamp steps recorded in a DRG neuron (26.2 μm diameter) in the absence (control; left) and presence of 100 μM baclofen (right). Broken line indicates 0 mV, black indicates the rheobase and red 2x rheobase in each condition for both membrane potential and current. **D** Bar graphs and scatter plots of the effects of 100 μM baclofen on resting membrane potential (RMP), p = 0.0177 **(i)**, input resistance (R_i_), p = 0.035 **(ii)**, rheobase, p = 0.0002 **(iii),** and number of APs, p = 0.0474 **(iv)** in response to 500 ms depolarizing current step in small to medium diameter neurons of adult mice DRG. Data represented as mean ± SEM; Paired t-test. Number of experiments is given in parentheses.

DRG neuronal excitability in response to depolarizing current pulses was greatly supressed in >two-thirds of neurons (21/31) upon application of baclofen (Fig. 5A). Baclofen (100 μM) hyperpolarized the resting membrane potential by ~5 mV from −59 ± 2 mV to −64 ± 2 mV (n = 18; p = 0.02) (Fig. 4C, Di, Table 2) and reduced the input resistance of DRG neurons approximately two-fold from 741.0 ± 189.5 MΩ (control) to 417.2 ± 99.4 MΩ (n = 10; p = 0.008) (Fig. 4Dii, Table 2). Baclofen also significantly increased the rheobase ~two-fold from 321 ± 88 pA to 620 ± 115 pA (n = 21; p = 0.0002) (Fig. 4Diii, Table 2) and attenuated the number of action potentials more than half in response to a 50 pA depolarizing current step (n = 21; p = 0.047) (Fig. 4Div, Table 2). These results are consistent with a reduction in DRG neuronal excitability in the presence of both Vc1.1 and baclofen.

**Figure 5.**
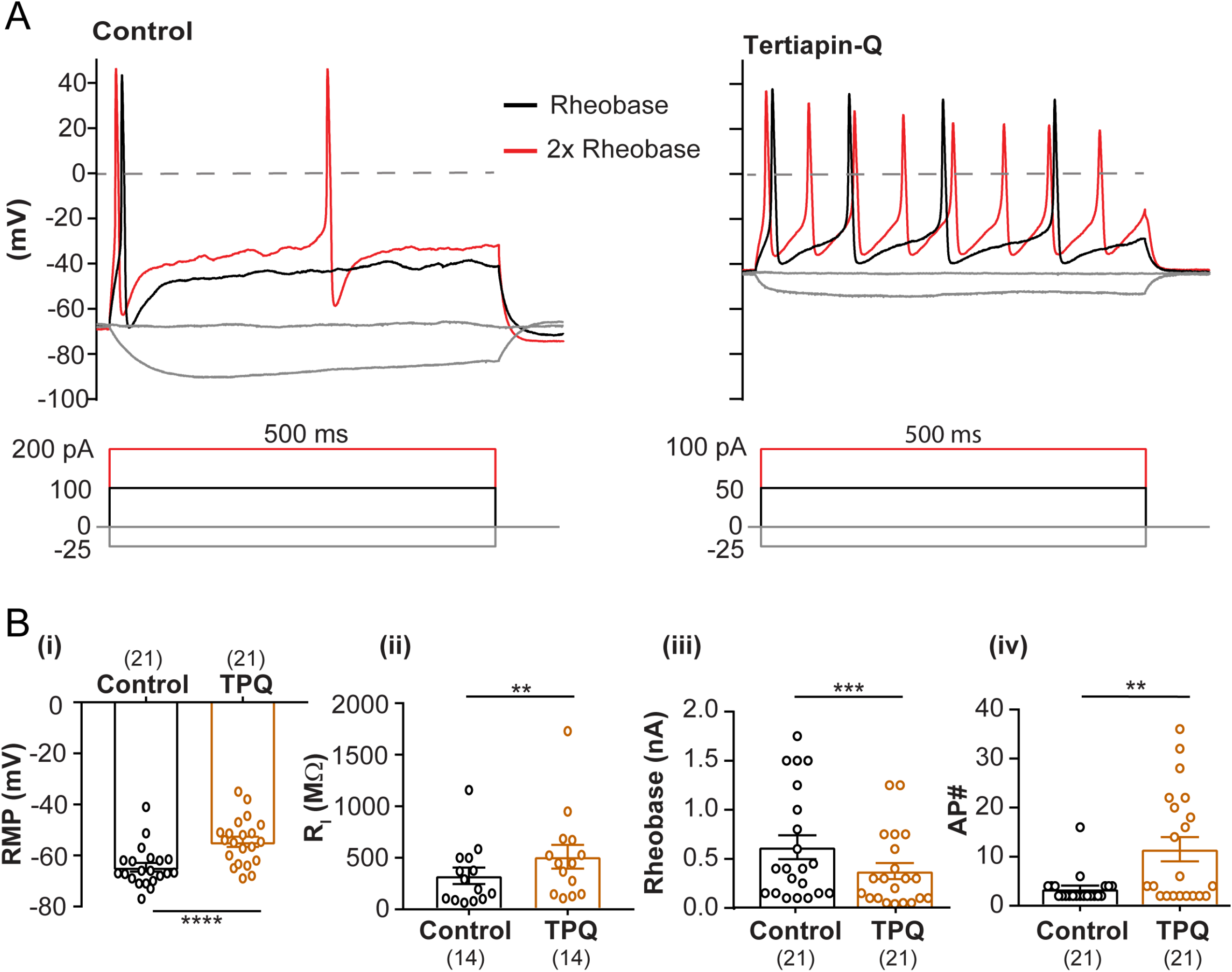
Tertiapin-Q, a GIRK channel inhibitor, enhances excitability in mouse DRG neurons and depolarizes the resting membrane potential. **A** Representative voltage responses to current clamp steps recorded in a DRG neuron (22.9 μm diameter) in the absence (control; left) and presence of 500 nM TPQ (right). Broken line indicates 0 mV, black indicates the rheobase and red indicates 2x rheobase for both membrane potential and current. **B** Bar graphs and scatter plots of the effects 500 nM Tertiapin-Q on resting membrane potential (RMP), p < 0.0001 **(i)**, input resistance (R_i_), p = 0.0354 **(ii)**, rheobase, p = 0.0003 **(iii)**, and number of APs, p = 0.0011 **(iv)** in response to 1 sec depolarizing current step in small to medium diameter (<30 μm) adult mice DRG neurons. Data represent mean ± SEM values; Paired t-test. Number of experiments is given in parentheses.

In contrast, spontaneous action potential firing was associated with a ~10 mV depolarization of the resting membrane potential from −65 ± 2 mV (control) to −55 ± 1 mV (n = 21; p < 0.0001) in the presence of 500 nM Tertiapin-Q (Fig. 5A, B; Table 2). Exposure of DRG neurons to Tertiapin-Q (500 nM) increased the input resistance more than ~1.5-fold from 326.0 ± 79.7 MΩ to 510.7 ± 114.7 MΩ (n = 14; p = 0.008) (Fig. 5C, Table 2). The rheobase was significantly reduced ~two-fold from 619 ± 122 pA to 376 ± 82 pA (n = 21; p = 0.0003) in the presence of 500 nM Tertiapin-Q (Fig. 5D, Table 2), and the number of action potentials elicited in response to a 25 pA depolarizing current step was increased more than three-fold (n = 21; p = 0.0011) (Fig. 5E, Table 2). In contrast, application of Vc1.1 in the presence of the GABA_B_R antagonist CGP 55845 did not alter passive or active membrane properties of mouse DRG neurons (n = 7; Fig. EV4A, B). Taken together, the results tabulated in Table 2 suggests that Vc1.1 and baclofen potentiation of GIRK-mediated K^+^ currents via GABA_B_R activation acts to dampen DRG neuronal excitability, and thereby contribute to the reported anti-nociceptive activities in animal models of mechanical allodynia and chronic pain (Klimis *et al*, 2011; Castro *et al*, 2017).

## Discussion

The major outcome of this study is the demonstration that analgesic α-conotoxin Vc1.1 modulates both GIRK1/2 and GIRK2 K^+^ currents through G protein-coupled GABA_B_R in HEK293T cells and mouse DRG neurons. Vc1.1 and baclofen potentiate recombinant GIRK1/2 and GIRK2 K^+^ currents only when co-expressed with GABA_B_R subunits R1 and R2. Inhibition of Vc1.1 potentiation of GIRK1/2 I_Kir_ by the selective GABA_B_R antagonist, CGP 55845, and the lack of effect of Vc1.1 on GIRK-mediated I_Kir_ in the absence of GABA_B_R expression, supports GABA_B_R as the primary target and not the GIRK channel itself. Our study also provides immunocytochemical support for the expression of GIRK channel protein in mouse DRG neurons.

GABA_B_ receptor biology and pharmacology in pain processing has been studied extensively and the modulation of HVA N-type calcium channels and/or GIRK channels by GABA_B_R agonists, GABA and baclofen, is a recognized mechanism of action (Malcangio, 2018; Benke, 2020). Our findings propose another plausible mechanism of action through which analgesic α-conotoxin Vc1.1 regulates membrane excitability in sensory neurons. Disruption of GABA_B_R-GIRK channel association in recombinant GABA_B_R/GIRK complexes HEK293 cells and native DRG neurons precludes I_Kir_ potentiation by baclofen and Vc1.1, consistent with previous studies supporting GIRK’s involvement in anti-nociception in models of neuropathic pain (Ippolito *et al*, 2005; Marker *et al*, 2005) and via satellite ganglion cells of the trigeminal ganglia (Takeda *et al*, 2015).

### Proposed mechanism underlying Vc1.1 activation of GABA_B_R

PTX catalyzes the ADP-ribosylation of the α_i_ subunits of the heterotrimeric G protein thereby precluding the interaction between G proteins and GPCRs. The involvement of the PTX-sensitive G protein, Gα_i/o_, in GABA_B_R modulation of GIRK channels by Vc1.1 was demonstrated by (1) pre-treatment of cells expressing either native or recombinant GABA_B_R/GIRK with PTX, and (2) intracellular application of the non-hydrolysable GDP analog, GDP-β-S, to inhibit of G protein activation. Both PTX and GDP-β-S abolished baclofen and Vc1.1 potentiation of GIRK1/2 channels. Furthermore, the observed inhibition of GIRK1/2 I_Kir_ potentiation by Vc1.1 and baclofen in the presence of the Gβγ scavenger, GRK2i, is consistent with Gβγ interaction with GIRK1/2 channels. GPCR-mediated activation of GIRK channel signaling involving Gα_i/o_ and Gβγ binding to the GIRK channel resulting I_Kir_ potentiation has been reported previously (Padgett & Slesinger, 2010; Wang *et al*, 2016; Kano *et al*, 2019). Fluorescence resonance energy transfer (FRET) studies show that upon GPCR activation Gβγ preferentially binds to the GIRK channel whereas Gα binds to GABA_B_R (Fowler *et al*, 2007; Laviv *et al*, 2011; Richard-Lalonde *et al*, 2013). Agonist dependent and GPCR-mediated activation of GIRK channels is proposed to occur via direct Gβγ subunit interaction with the channel in a membrane-delimited fashion (Lujan *et al*, 2014; Yudin & Rohacs, 2018). The recruitment of Gβγ by the GIRK channel is essential in GPCR-mediated potentiation, whereas other prototypical agonists that activate GIRK channels in the absence of a GPCR or in a G protein-independent manner, do not require the Gβγ heterodimer (Jelacic *et al*, 2000; Wydeven *et al*, 2014). Accordingly, our results provide evidence that Vc1.1 targeting of GABA_B_R modulates GIRK channels via activation of Gα_i/o_ protein and Gβγ signaling as proposed in Fig. 6.

**Figure 6.**
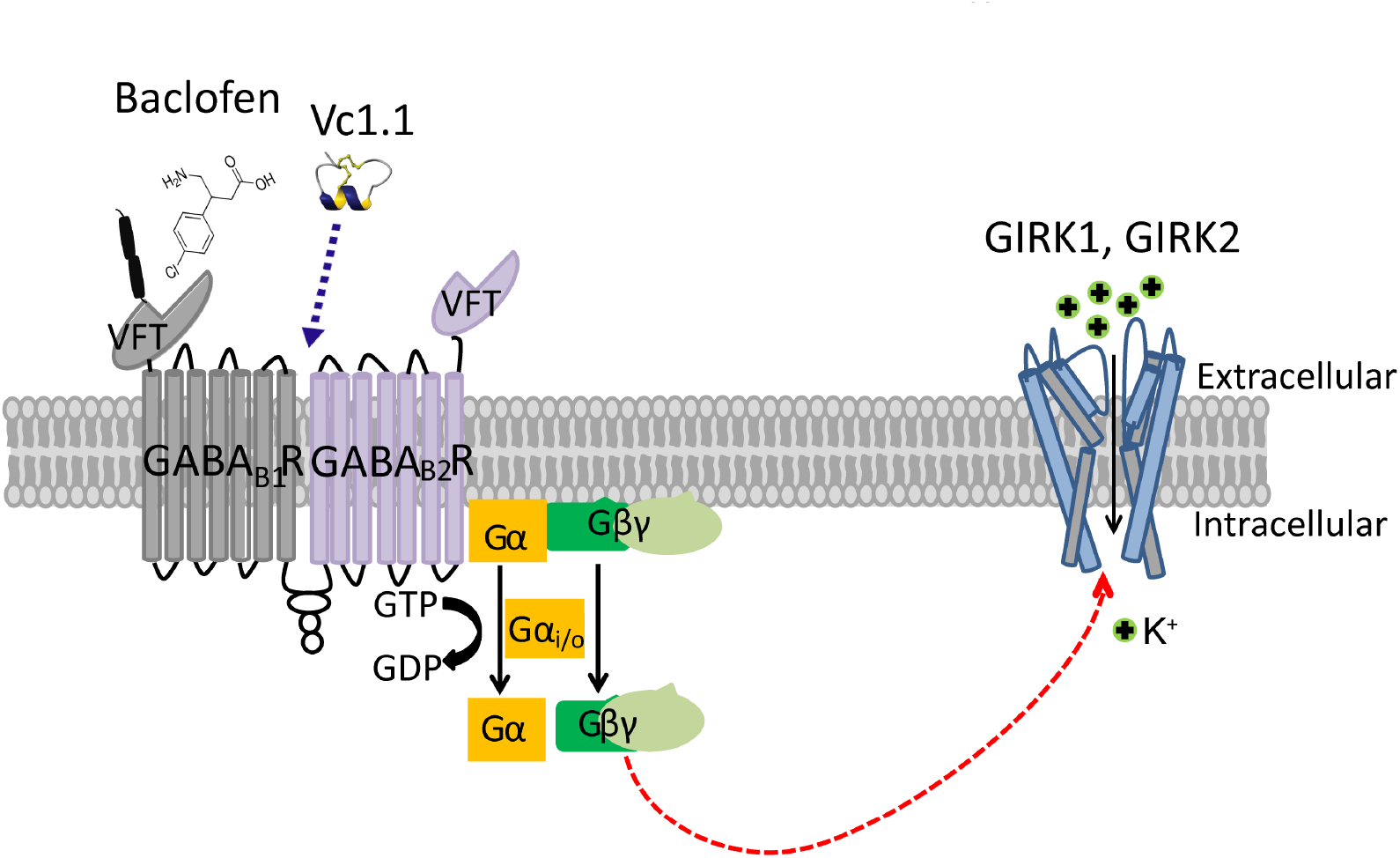
Schematic representation of the GABA_B1/2_ subunits of GABA_B_R and its downstream effector Vc1.1 transducing signals to GIRK channels. Allosteric binding of Vc1.1 to GABA_B_R activates heterotrimeric G protein. Dissociation of Gα_i/o_ and heterodimer Gβγ (red line) of G protein activates heteromeric GIRK1/2 channel. Similarly, baclofen binding to orthosteric binding site, Venus Fly Trap (VFT) of GABA_B1_ subunit of GABA_B_R can also potentiate GIRK channel via dissociation of Gα_i/o_ and Gβγ heterodimer of G protein.

α-Conotoxin Vc1.1 has been shown to act as an allosteric agonist as it does not bind to the canonical ligand binding site of GABA_B_R (Huynh *et al*, 2015), nevertheless, its precise binding site on the GABA_B_R remains to be identified (Sadeghi *et al*, 2017). In the present study, we show that Vc1.1 potentiates GIRK1/2 channels albeit with 10- to 100-fold lower potency than for inhibition of human and rat Ca_v_2.2 channels (Callaghan & Adams, 2010; Huynh *et al*, 2015; Hone *et al*, 2018). In a comparable manner, it was shown that even though GIRK and Cav2.2 channels are regulated by GABA_B_R by an analogous mechanism, the EC_50_ for GIRK potentiation by baclofen is ~100-fold higher than for inhibition of HVA Ca^2+^ channels in rat supraoptic neurons (Harayama *et al*, 2014).

A durable macromolecular complex between GIRK-G protein-GABA_B_R upon agonist binding (Padgett & Slesinger, 2010) that occurs through collision coupling (rather than a pre-existing complex) has been proposed (Kahanovitch *et al*, 2017). It is conceivable that Vc1.1 targets the complex activating Gα_i/o_ proteins and Gβγ via GABA_B_R to reversibly potentiate GIRK channel signaling. This is consistent with previous studies whereby agonist stimulation of Gα_i/o_-coupled GPCRs results in increased concentration of Gβγ heterodimers available to activate GIRK channels whereas the Gα_i/o_ subunit enhances G protein-GPCR association in GPCR-GIRK complexes (Touhara & McKinnon, 2018).

### Expression and modulation of GIRK channels in sensory neurons

GIRK channels are expressed in neurons of several regions of the central nervous system including the spinal cord (Ponce *et al*, 1996; Marker *et al*, 2006; Lujan & Aguado, 2015) supporting their role in modulating several neurological disorders or disease conditions (Mayfield *et al*, 2015; Rifkin *et al*, 2017). Functional coupling to GABA_B_R has been described in rat (Gao et al, 2007) and human DRG neurons (Castro *et al*, 2017), yet the function of GIRK channels in mouse peripheral neurons has remained largely unexplored.

Functional recordings of phosphatidylinositol 4,5-bisphosphate [PtdIns(4,5)P2/PIP2] activated GIRK currents in postnatal day zero (P0) mouse DRG neurons in response to neurotrophin-mediated cleavage of p75(NTR) have been reported (Coulson *et al*, 2008). Here, we show functional coupling between GIRK1/2 (and GIRK2) channels and GABA_B_R in mouse DRG neurons and heterologous expression systems evidenced by the potentiating actions of baclofen and Vc1.1 on I_Kir_. This is supported by co-localization of GIRK1, GIRK2, and GABA_B_R2 subunits in overlapping populations of small and medium diameter neurons of adult mouse DRG (Fig. 3A, B).

In voltage clamp experiments, approximately one-third of DRG neurons responded to Vc1.1 which is similar to the proportion of mouse DRG neurons in which the μ-opioid receptor agonist DAMGO elicited I_Kir_-like currents reported by Saloman *et al*. (2016). Similar to previous studies of baclofen treated rat trigeminal ganglion and reticular thalamic neurons (Takeda *et al*, 2004; Cain *et al*, 2017), baclofen and Vc1.1 hyperpolarized the resting membrane potential, increased the firing threshold (rheobase) and reduced action potential firing in the mouse DRG neurons analyzed here. The GABA_B_R antagonist CGP55845 abrogated potentiation of I_Kir_ and the neuroexcitability dampening effects of Vc1.1 whereas Ba^2+^ and the GIRK channel inhibitor Tertiapin-Q enhanced excitability and antagonized the potentiation of I_Kir_ by baclofen and Vc1.1 in mouse DRG neuron. Thus far, our findings are consistent with GABA_B_R modulation of GIRK channels and the concomitant reduction in DRG neuronal excitability. Tertiapin-Q antagonized the effects of Vc1.1 and baclofen on neuronal excitability in a manner consistent with results of our study of human GABA_B_R and GIRK1/2 channels expressed in HEK293T cells. Hence, GIRK channels function as downstream effectors of GABA_B_R contributing to the anti-nociceptive activity of baclofen and Vc1.1.

In rat pyramidal neurons, GIRK channel density is higher in dendrites than in the soma (Takigawa & Alzheimer, 1999; Chen & Johnston, 2005) perhaps posing a limitation to our work as it relies on somatic DRG neuron I_Kir_ recordings (within 24 hours of acute dissociation). The relatively low I_Kir_ density in adult mouse DRG neurons implies that most processes and the dendritic pool of GIRK channels that may be associated are lost upon isolation. Future efforts will be directed to determine the relative abundance of GIRK channel isoforms at the mouse peripheral nerve terminals.

#### Neuroexcitability and implications for GIRK channel

Analgesic α-conotoxins inhibit α9α10 nAChRs directly and HVA calcium (Cav2.2 and Cav2.3) channels via GABA_B_R activation (Sadeghi *et al*, 2017; Kennedy *et al*, 2020). The GABA_B_R is abundantly expressed in the somatosensory (afferent) nervous system and its role in mitigating mechanical allodynia and chronic visceral hypersensitivity in animal models of neuropathic and visceral pain, respectively, has been demonstrated (Castro *et al*, 2017; Loeza-Alcocer *et al*, 2019). Furthermore, α-conotoxin Vc1.1 has been shown to reduce excitability in human DRG neurons via a GABA_B_R-mediated mechanism. In the present study, we demonstrate that analgesic α-conotoxins Vc1.1, RgIA and PeIA potentiate heteromeric and homomeric GIRK currents via GABA_B_R. However, the GABA_B_R agonists, baclofen and GABA, tested at maximally effective concentrations, were more efficacious than the analgesic α-conotoxins, Vc1.1, RgIA, and PeIA in potentiating of GIRK-mediated I_Kir_.

Hyper-excitability and ectopic firing are characteristic sensory neuron responses to nerve injury and chronic pain (Amir *et al*, 2005; Berta *et al*, 2017). Control of cell-surface expression and/or the biophysical properties of ion channels in sensory neurons is central to membrane excitability such that the activation of GIRK channels hyperpolarizes the neuron thus reducing excitability. Our findings are consistent with the functional expression of GIRK channels in mouse DRG neurons and their involvement in GABA_B_R-mediated anti-nociceptive activity in response to baclofen and α-contotoxin Vc1.1. The modulation of HVA N-type (Cav2.2) calcium channels and/or GIRK channels by GABA_B_R agonists, GABA and baclofen, is recognized as analgesic mechanism of action. A new scenario involving diverse signaling mechanism(s) underlying α-conotoxin Vc1.1’s analgesic effect is emerging. Synergistic action of Vc1.1 to inhibit Cav2.2/2.3 and potentiate GIRK channels reducing neuronal excitability likely contributes to its anti-nociceptive activity. Lastly, the precise molecular details of GIRK I_Kir_ potentiation via GABA_B_R activation remains to be elucidated, therefore, further studies of sensory neuron GIRK channels and their coupling to GPCRs will follow.

## Materials and Methods

### Antibodies and pharmacological agents

Primary antibodies anti-GIRK1 and anti-GIRK2 were purchased from Alomone Labs (Jerusalem, Israel). The primary antibodies were rabbit polyclonal GIRK1 (P63250, intracellular, C-terminus), guinea pig polyclonal GIRK2 (P48542, intracellular, C-terminus), rabbit polyclonal GIRK2 (P348542, intracellular, C-terminus) and rabbit monoclonal GABA_B_R2 (Abcam, ab75838). The secondary antibodies (Abcam Cambridge, MA USA) were Alexa Fluor 647-conjugated polyclonal donkey anti-rabbit IgG antibody (ab150075), Alexa Fluor 488-conjugated polyclonal goat anti-guinea pig IgG antibody (ab150185) and Alexa Fluor 488-conjugated polyclonal goat anti-rabbit IgG antibody (ab150073). DAPI (4′,6-diamidino-2-phenylindole; Sigma-Aldrich, North Ryde, NSW, Australia) was used for nuclear staining of the neurons.

α-Conotoxins Vc1.1, PeIA, Rg1A, and ImI were synthesized as described previously (*10, 11, 15*) and kindly provided by Dr Richard Clark (University of Queensland, Australia). (±)-β-(Aminomethyl)-4-chlorobenzenepropanoic acid (Baclofen), γ-aminobutyric acid (GABA), guanosine 5′-[β-thio]diphosphate trilithium salt (GDP-β-S), guanosine 5′-[γ-thio]triphosphate tetralithium salt (GTP-γ-S) and *Pertussis* toxin (PTX) were purchased from Sigma-Aldrich (St. Louis, MO), CGP 55845 ((2S)-3-[[(1S)-1-(3,4-Dichlorophenyl)ethyl]amino-2-hydroxypropyl] (phenylmethyl) phosphinic acid hydrochloride) and GRK2i were purchased from Tocris Bioscience (Bristol, UK). Tertiapin-Q, a potent inhibitor of inward rectifier K^+^ (Kir) channels, was purchased from Abcam (Cambridge, UK). All drugs were dissolved in distilled H_2_O to prepare to their appropriate stock concentration other than CGP 55845 which was dissolved in DMSO. Compounds were then diluted in extracellular high K^+^ (20 mM). The final concentration of DMSO did not exceed 0.01%.

### Cell culture, transfection and clones

HEK293T cells expressing the SV40 large T antigen (HEK293T, ATCC, USA) were cultured in Dulbecco’s modified Eagle’s medium (DMEM, ThermoFisher Scientific, Scoresby, VIC, Australia), supplemented with 10% heat inactivated foetal bovine serum (FBS, Bovigen, Australia), 1% penicillin and streptomycin (Pen/Strep) and 1% GlutaMAX (ThermoFisher Scientific, Australia). Cells were incubated at 37°C in 5% CO_2_ in a humidified incubator and passaged when ~80% confluent following the procedures described previously.

HEK293T cells were transiently co-transfected with plasmid cDNAs encoding human GIRK 1 (KCNJ3 or Kir3.1) and/or GIRK 2 (KCNJ6 or Kir3.2) (both pcDNA3 based and were kindly provided by Dr Paul Slesinger, Mt Sinai, New York, USA) and human GABA_B1_ and GABA_B2_ subunits (OriGene Technologies, Inc, Rockville, MD USA) using Lipofectamine 2000 (ThermoFisher Scientific, Australia). Cells were seeded in 12-well plates the day before transfection and 2 μg of each plasmid DNA with 0.2 μg of green fluorescent protein (GFP) were transfected following the manufacturer’s protocol of Lipofectamine. Cells were then replated on 12 mm glass coverslips 48-72 hours after transfection for subsequent patch clamp studies.

### Whole-cell patch clamp of HEK293T cells

Transiently transfected HEK293T cells were recorded 48-72 hours post transfection using whole-cell patch clamp methods. An external bath solution containing high K^+^ was used to record GIRK K^+^ currents that included (in mM): 120 NaCl, 20 KCl, 2 CaCl_2_, 1 MgCl_2_, 10 HEPES and 10 Glucose, pH 7.4 with NaOH (~320 mOsmol.kg^−1^). Borosilicate glass electrodes (World Precision Instruments, Sarasota, FL USA) with an access resistance of 3-5 MΩ were filled with internal solution containing (in mM): 130 KCl, 20 NaCl, 5 EGTA (ethylene glycol-bis(β-aminoethyl ether)-N,N,N′,N′-tetra acetic acid), 5.46 MgCl_2,_ 10 HEPES, 5 MgATP, 0.2 Na-GTP, pH 7.2 with KOH (~300 mOsmol.kg^−1^). Membrane currents were recorded using a MultiClamp 700B amplifier and digitized with a Digidata 1440A (Molecular Devices, San Jose, CA USA) with series resistance (<10 MΩ) and cell capacitance compensated by ~80% avoiding current oscillation. All membrane currents were sampled at 10 kHz and filtered at 1 kHz.

GIRK K^+^ currents were recorded by applying a voltage ramp protocol from −100 mV to +40 mV from a holding potential of −40 mV, at a frequency of 0.1 Hz. Modulation of GIRK K^+^ currents by bath application of various compounds was measured at −100 mV. All experiments were carried out at room temperature (21-23°C).

### Isolation and culture of rodent dorsal root ganglion (DRG)

All animal procedures were conducted in accordance with the University of Wollongong Animal Ethics Committee (AE16/10) guidelines and regulations. Adult C57BL/6 mice (8-10 weeks old) were purchased from Australian BioResources (Moss Vale, NSW, Australia) and housed in individually ventilated cages with a 12 h light/dark cycle; food pellets and water were available *ad libitum*.

Mice were euthanized by isoflurane inhalation followed by rapid decapitation. A laminectomy exposed the DRGs in the thoracic and lumbar regions. DRGs were harvested and transferred to ice cold (4°C) Hanks Buffered Saline Solution (HBSS), free of Ca^2+^ and Mg^2+^. DRGs were then trimmed, removing central and peripheral nerve processes, digested in HBSS containing Collagenase type II (3 mg/ml; Worthington Biomedical Corp., Lakewood, NJ, USA) and Dispase (4 mg/ml; GIBCO, Australia). The DRGs were incubated in this enzyme mix at 37°C in 5% CO_2_ in a humidified incubator for 40 min. The ganglia were then rinsed three to four times in warm (37°C) F12/Glutamax (Invitrogen) media supplemented with 10% heat-inactivated foetal bovine serum (FBS; GIBCO™, ThermoFisher Scientific, Australia) and 1% penicillin/streptomycin. The ganglia were dispersed by mechanical trituration with progressively smaller fire polished glass Pasteur Pipettes. The supernatant was filtered through a 160 μm nylon mesh (Millipore Australia Pty Ltd, North Ryde, NSW) to remove undigested material. The dissociated DRG neurons were then plated onto poly-D-lysine coated 12 mm cover glass (Sigma-Aldrich, Australia). Neurons were left to attach for ~3 hours at 37°C after which media was added and then incubated overnight and used within 24 hrs.

### Current and voltage clamp recording of native GIRK currents

Current clamp recordings of DRG neurons were typically carried out 24 hours after dissociation in a bath solution containing (in mM): 140 NaCl, 4 KCl, 2 CaCl_2_, 1 MgCl_2_, 10 HEPES and 10 Glucose, pH 7.4 with NaOH (~320 mOsmol.kg^−1^). Borosilicate fire polished patch pipettes (World Precision Instruments) with a resistance of 2-4 MΩ were filled with standard internal solution containing (in mM): 130 KCl, 20 NaCl, 5 EGTA (ethylene glycol-bis(β-aminoethyl ether)-N,N,N′,N′-tetraacetic acid), 5.46 MgCl_2,_ 10 HEPES, 5 Mg-ATP, 0.2 Na-GTP, pH 7.2 with KOH (~300 mOsmol.kg^−1^). Action potentials were generated by series of depolarizing current steps of 500 ms duration in small to medium diameter (<30 μm) DRG neurons that had a resting membrane potential (RMP) of > −40 mV. Whole-cell configuration was obtained in voltage clamp mode before proceeding to current clamp recording. Neuronal excitability was measured every 5 s using a constant amplitude small depolarizing current pulses.

Whole-cell GIRK channel K^+^ currents were recorded from isolated DRG neurons superfused with high K^+^ extracellular solution containing (in mM): 120 NaCl, 20 KCl, 2 CaCl_2_, 1 MgCl_2_, 10 HEPES and 10 Glucose, pH 7.4 with NaOH (~320 mOsmol.kg^−1^). Patch pipettes were filled with the same intracellular solution as above. Currents were elicited using a similar voltage protocol to that used for HEK293 cells (ramp from −100 mV to +40 mV, at a frequency of 0.1 Hz, from a holding potential of −40 mV and sampled at 10 kHz). Recordings were electronically compensated for cell capacitance and series resistance to ~80%. All solutions including the pharmacological agents were superfused using a Peristaltic pump at an exchange speed of 1 ml/min. The volume of the experimental chamber was ~500 μl. All experiments were carried out at room temperature (21-23°C).

### Immunocytochemistry and confocal microscopy

Dissociated DRG neurons plated on poly-D-lysine coated glass coverslips were washed twice with HBSS and fixed using Zamboni’s fixative solution containing 1.6% formaldehyde (Australian Biostain Pty Ltd, Traralgon, VIC, Australia) for 15 min. Cells were permeabilized with 0.1% Triton X-100 for 10 min. Non-specific antibody binding was reduced by incubating the cells for 1 hour in HBSS based blocking solution containing 5% goat serum, 5% BSA and 0.1% Triton X-100. All primary and secondary antibodies were diluted in blocking solution to the working concentration. For DRG neuron double staining of GIRK1 and GIRK2 channels, both anti-rabbit polyclonal GIRK1 (1:100 dilution) and anti-guinea pig polyclonal GIRK2 (1:100 dilution) antibodies were added simultaneously and incubated at 4°C overnight. Following three 5 min washes with blocking solution, the samples were incubated with the secondary antibody, a donkey anti-rabbit IgG (1:500) 1 hr, in the dark, at room temperature (RT). Triplicate washes with blocking solution preceded addition of the goat anti-guinea pig IgG secondary antibody (1:500) which again was incubated 1 hr at RT, in the dark. For GIRK2 and GABA_B_R2 double staining, a mixture of the primary antibodies rabbit polyclonal GIRK2 (1:100) and rabbit monoclonal GABA_B_R2 (1:400) were incubated at 4°C overnight, washed and followed by sequential incubations with donkey anti-rabbit IgG secondary antibody (1:500, 1 hr, RT), wash and goat anti-rabbit IgG secondary (1:500, 1 hr, RT). Control experiments for single and double staining procedures were performed by omitting the primary antibodies. Nuclei were counterstained with DAPI (1:5000, 5 min, RT), washed, covered with mounting media (Dako North America Inc., Carpinteria, CA, USA), sealed and stored at 4°C. Images were recorded with a Leica SP8 confocal microscope using 40x oil immersion objective. Images were analysed using ImageJ (Java, NIH) and LAS X softwares (Leica, Macquarie Park, NSW, Australia).

### Data analysis and statistics

Data analysis used Clampfit 10.7 (Molecular Devices, San Jose, CA, USA) and GraphPad Prism 7 (GraphPad Software, San Diego, CA, USA) software. In voltage clamp experiments, inward rectifying K^+^ currents (I_Kir_) were analysed by measuring the steady-state current in high K^+^ solution designated basal I_Kir_ (I_CONTROL_, I_CTR_), peak potentiation by agonists (I_COMPOUND_, I_COMP_) and I_Kir_ density (pA/pF) determined by dividing peak current amplitude by cell capacitance (Cm). The Ba^2+^-sensitive current amplitude was determined in the presence of 1 mM Ba^2+^ (in high K^+^ bath solution) and subtracting the current before and after application of the modulators to evaluate the effects of GABA_B_R-active modulators on the K^+^ current.

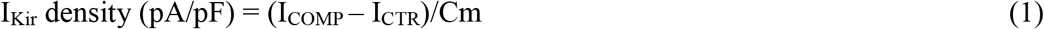

Concentration-response relationship for Vc1.1 was determined using equation (1) from 4-5 individual experiments for each concentration tested on GIRK1/2 channels. The relationship was fitted with the Hill equation following:

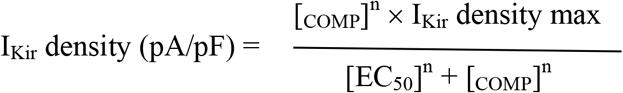

where *n* is the Hill coefficient, EC_50_ is the half-maximal response.

In current clamp experiments, all DRG neurons accepted for analysis had a RMP more negative than −40 mV. The whole-cell input resistance (R_i_) was measured in the absence and presence of various modulators by calculating the slope of the linear fit of hyperpolarizing responses to current steps from −5 to −40 pA in 5 pA increments. Rheobase threshold was measured by applying a series of depolarizing current steps (500 ms) in 5 pA increments to determine which first evoked action potential discharge. The number of action potentials was counted at a current injection of 2x rheobase in both control conditions and in the presence of modulators or pharmacological agents.

Statistical significance was determined between two groups by using paired t-test or Student’s t-test. Multiple group comparisons were done by one-way ANOVA with Tukey’s test. All results are presented as mean ± SEM and n, number of observations. Statistical significance is shown by asterisk *indicating p < 0.05, **indicating p < 0.005 and ***indicating p < 0.001 and ****indicating p < 0.0001.

## Data Availability

This study includes no data deposited in external repositories.

## Acknowledgements

We thank Dr Alexander (Sandy) Harper, University of Dundee, for his insightful comments on a draft manuscript. A.R.B. was supported by a University of Wollongong Postgraduate Research Scholarship. This work was supported by an Australian National Health and Medical Research Council (NHMRC) Program Grant [APP1072113] to D.J.A

## Author contributions

D.J.A. conceived and supervised the project; D.J.A., J.R.M. and R.K.F-U. designed research; A.R.B. performed research; A.R.B, J.R.M, R.K.F-U. and D.J.A. analyzed the data; A.R.B, J.R.M, R.K.F-U. and D.J.A. wrote the paper.

## Conflict of interest

The authors declare no competing financial interests.

## Expanded View Figure Legends

**Figure EV1.**
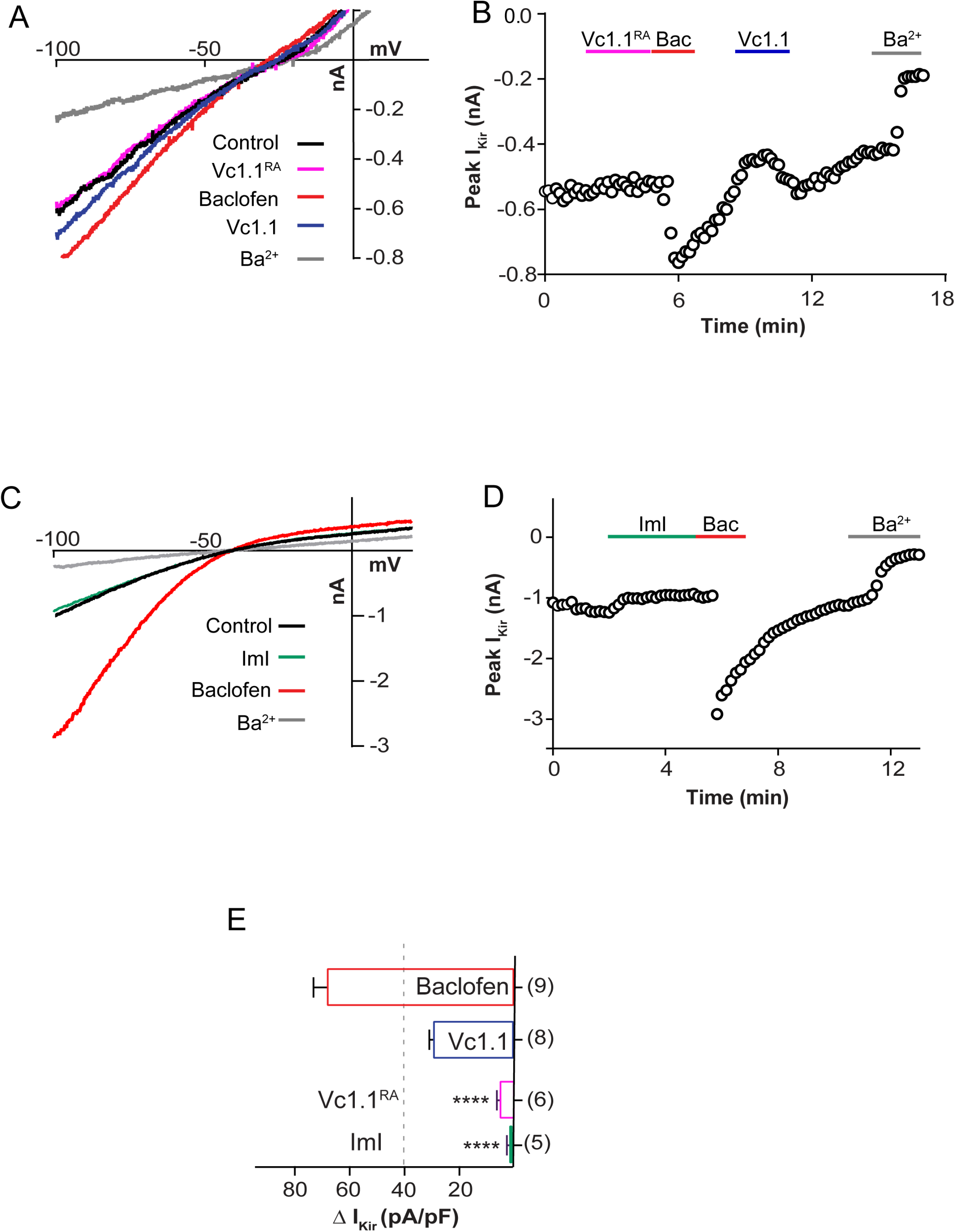
α-Conotoxin Vc1.1 activity abolished upon reducing the disulfide bond with dithiothreitol (DTT) and the lack of effect of α-conotoxin ImI on heteromeric GIRK1/2 channels co-expressed with GABA_B_R. **A** Representative current-voltage relationship elicited from HEK293 cells co-expressing GABA_B_R and GIRK1/2 channels in the absence (Control, black) and presence of 1 μM Vc1.1 reduced by 1 mM DTT/reduced Vc1.1 (Vc1.1^RA^) (pink), 100 μM baclofen (red), 1 μM wild-type Vc1.1 (blue) or 1 mM Ba^2+^ (grey). **B** GIRK1/2-mediated K^+^ current amplitude measured at −100 mV plotted as a function of time in the presence of sequential application of reduced Vc1.1 (DTT treatment/Vc1.1^RA^), baclofen, globular Vc1.1 and Ba^2+^. **C** Representative current-voltage relationships obtained from HEK293T cells co-expressing GABA_B_R and GIRK1/2 channels in the absence (Control, black) and presence of 1 μM ImI (light blue), 1 μM Vc1.1 (blue), 100 μM baclofen (red) or 1 mM Ba^2+^ (grey). **D** Peak K^+^ current amplitude plotted as a function of time during sequential application of ImI, Vc1.1, baclofen, and Ba^2+^. **E** Quantification of the effect of reduced (linear) Vc1.1 and ImI compared to globular Vc1.1 and baclofen. Data represent mean ± SEM. Statistical significance, **** p < 0.0001; one-way ANOVA. Number of experiments is given in parentheses.

**Figure EV2.**
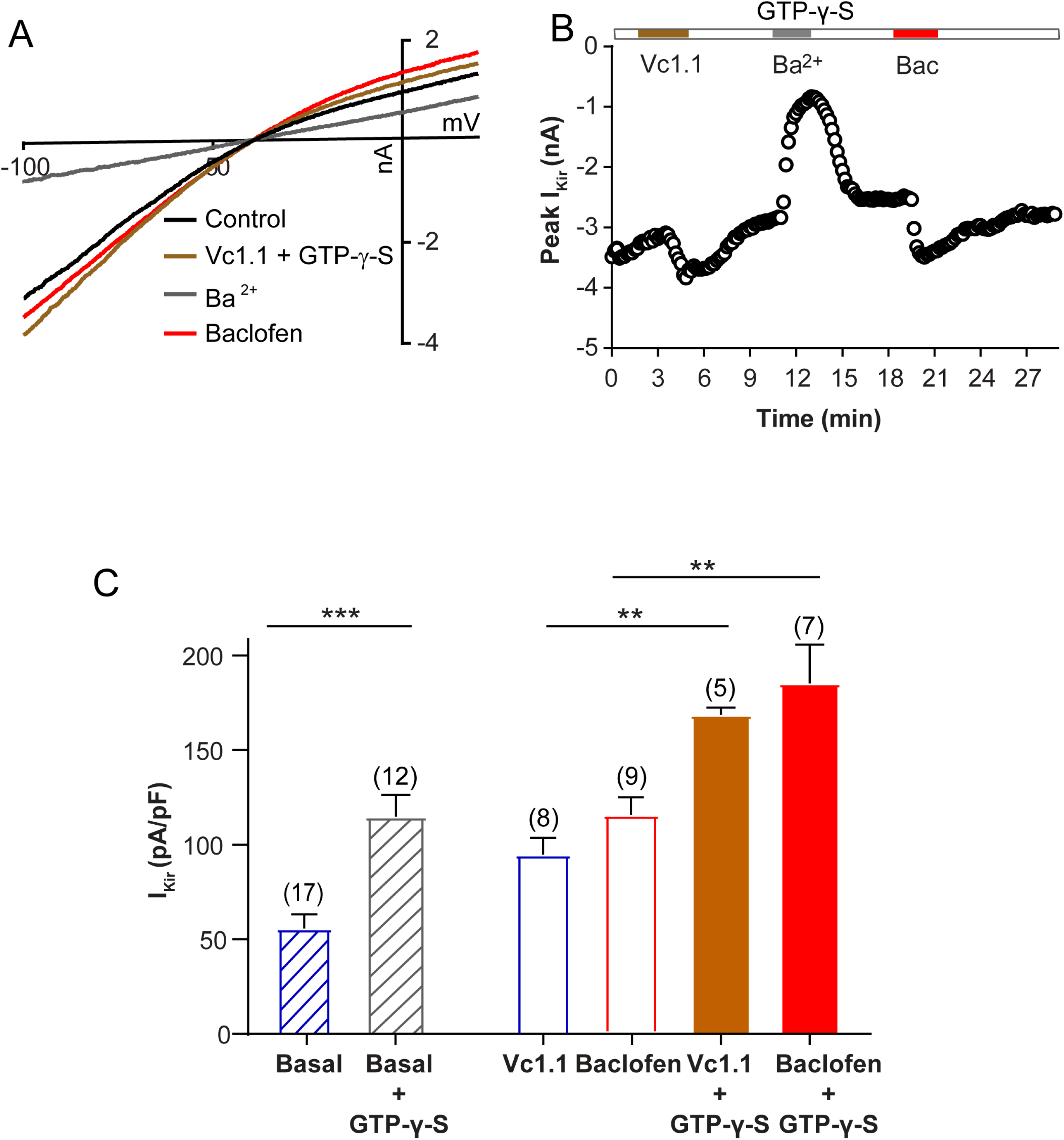
Heterotetrameric GIRK1/2 K^+^ currents enhanced by intracellular GTP-γ-S. **A** Current-voltage relationship obtained in HEK293T cells containing intracellular GTP-γ-S (500 μM) in the absence (basal) and presence of Vc1.1 (1 μM, brown), Ba^2+^ (1 mM, grey) and baclofen (100 μM, red). **B** Corresponding plot of peak I_Kir_ density as a function of time in HEK293T cell containing GTP-γ-S. **C** Bar graph comparing the effect of the absence (control, Vc1.1) and presence of GTP-γ-S (500 μM) on basal, Vc1.1 (1 μM) and baclofen (100 μM) induced peak I_Kir_ density. Data are expressed as mean ± SEM; statistical significance *** p = 0.004, ** p = 0.005 and p = 0.002 respectively; one-way ANOVA followed by Tukey’s post hoc test; number of experiments is given in parentheses.

**Figure EV3.**
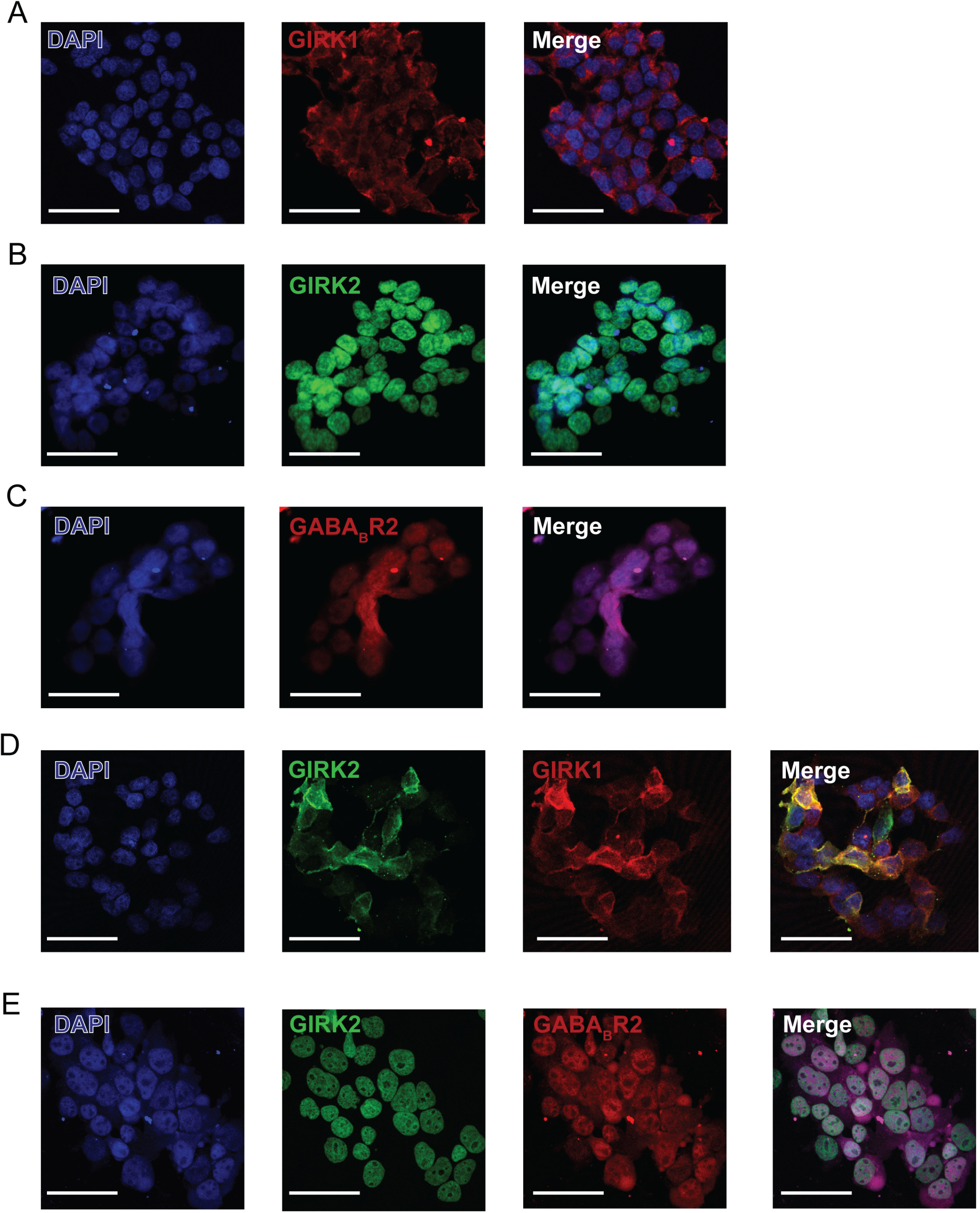
Fluorescence micrographs representing immunolabeling of GIRK 1, GIRK 2 and GABA_B_R2 alone and co-localization when over-expressed recombinantly in HEK293 cells. **A-C** GIRK1 (red), GIRK2 (green) and GABA_B_R2 (red) exhibited immunoreactivity in HEK293T cells and DAPI (blue) staining denotes the nucleus of the cells. **D-E** DAPI (blue), GIRK1 (red), GIRK2 (green) and GABA_B_R2 (red) immunoreactivity in HEK293T cells. The merged/overlay images depict co-localization of GIRK1 and GIRK2 or GIRK2 and GABA_B_R2 upon transfection. Scale bars: 100 μm.

**Figure EV4.**
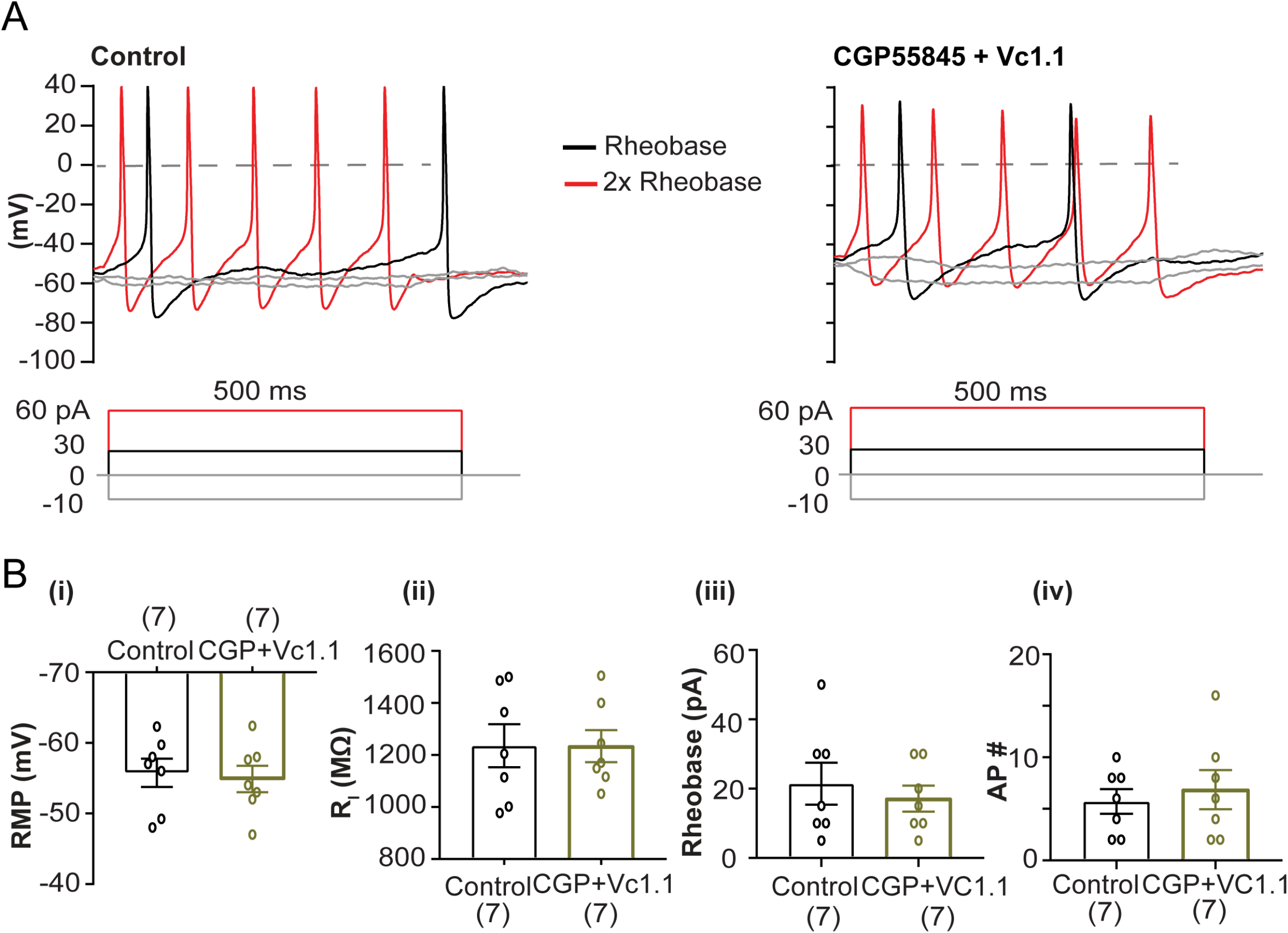
Vc1.1-induced excitability in mouse DRG neurons is antagonized by the selective GABA_B_R antagonist CGP55845. **A** Representative voltage responses to current clamp steps recorded in a DRG neuron (26 μm diameter) in the absence (control; left) and presence of CGP 55845 + Vc1.1 (1 μM) (right). Broken line indicates 0 mV, black indicates the rheobase and red indicates 2x rheobase for both membrane potential and current. **B** Bar graphs and scatter plots of the effects CGP 55845 + Vc1.1 on resting membrane potential (RMP), p = 0.4 **(i)**, input resistance (R_i_), p = 0.1 **(ii)**, rheobase, p = 0.36 **(iii)**, and number of APs, p = 0.39 **(iv)** in response to 500 ms depolarizing current step in small to medium diameter neurons of adult mouse DRG. Data represent mean ± SEM values; Paired t-test. Number of experiments is given in parentheses.

